# TOR signalling regulates epithelial cell shape transition in *Drosophila* oogenesis

**DOI:** 10.1101/2022.12.29.522192

**Authors:** Sudipta Halder, Gaurab Ghosh, Budhaditya Gayen, Mohit Prasad

## Abstract

Epithelial morphogenesis plays an important role in form generation, organ development, and maintenance of adult tissues in the metazoans. Given its wide implication, aberrant morphogenesis is linked to severe developmental defects and in a few instances also associated with tumorigenesis. Employing the model of *Drosophila* oogenesis, we have examined the role of the evolutionary conserved Target of Rapamycin (TOR) kinase pathway, a known regulator of cell growth and size in mediating shape transition of cuboidal cells to squamous epithelial fate. Utilizing genetic tools, immunohistochemistry, and live cell imaging, we demonstrate that TOR signaling is active and required for epithelial morphogenesis during *Drosophila* oogenesis. Further, loss of function analyses indicates that non-canonical TOR signaling functions through PAR-1 to mediate the removal of lateral cell adhesion molecule, Fasciclin 2, to allow proper squamous cell morphogenesis. In addition, we demonstrate the effect of TOR through PAR-1 on cell shape transition is mediated via the modulation of endocytosis. Overall, our data give novel insight into how TOR signaling mediates cell shape transition during epithelial morphogenesis in the metazoans.

## Introduction

The evolutionarily conserved Target of rapamycin (TOR) pathway plays an important role in coupling metabolism with cellular growth both in development and diseases. As most of the TOR-regulated processes are linked to cellular growth, the role of TOR in aiding expansive cell shape transformations hasn’t been examined so far.

The simple squamous epithelia are quite prevalent in the metazoan, known to line the cavities of the mammalian body, blood vessels and are also found in kidney glomeruli and the respiratory tract ^1^. Incidentally, quite a few of the squamous cell fates like those observed in the mature alveoli lining of lungs, the parietal epithelial cells of the glomerulus, or the stratum corneum of the mammalian skin are derived from cuboidal cells undergoing epithelial morphogenesis ^1–3^. As expected, these expansive epithelial cell shape transformations are assisted with changes in the cytoskeletal network and remodelling of the adherens junctions ^4–7^, however, the identity of factors that affect this change is unclear. As cuboidal to squamous shape transition is associated with cell growth, one of the open questions that still intrigues the field is how the shape and size of epithelial cells are regulated during development.

The Target of Rapamycin (TOR) is a serine/ threonine kinase and in response to diverse stimuli, it functions through two distinct complexes (TORC1 and TORC2) to mediate cellular metabolism through modulation of protein synthesis, mRNA stability, and/ or autophagy ^8–14^. By temporally modulating the cell cycle phases, TORC1 regulates cell size ^15, 16^, while TORC2 affects spatial growth by regulating glucose metabolism, actin organization and suppressing apoptosis ^17–20^. Though TOR signaling regulates cell size, it is not clear whether it has any role in morphogenesis where cell shape transitions are associated with expansive growth. Intriguingly, expansive cell shape transformations constitute the backbone of metazoan morphogenesis from aiding early gastrulation through potentiating organ development to mediating repair and regeneration in the adults. Of all the cell shape transitions, we specifically focussed on the transformation of cuboidal epithelial cells to squamous fate as it is conspicuous, frequently associated with metazoan development, and most importantly, is the least understood so far.

*Drosophila* oogenesis has emerged as a genetically tractable system to understand various aspects of metazoan epithelial morphogenesis including growth, fate specification, group cell movement, and epithelial cell shape transformations ^21–23^. Typically, a fly egg consists of 16 germline nurse cells enveloped by 750 cuboidal epithelial cells. As the egg grows in size with the deposition of the maternal components, the germline associated cuboidal follicle cells undergo conspicuous shape transition with the anteriorly localized cells (∼50) undergoing flattening and stretching to acquire squamous fate while the posterior cells elongate to take columnar fate ^24^. The follicle cells in the mid-region of the egg chamber migrate to form the dorsal appendages ^24^. As the cuboidal to squamous shape transition is significant, we focussed our attention on how the growth promoting molecules regulate the shape transition of anterior follicle cells (AFCs) from cuboidal to squamous fate.

By coupling morphometric analysis and mathematical modelling, it has been proposed that the growth of the germline nurse cell induces flattening and stretching through Notch-mediated disassembly of adherens junctions in the overlying cuboidal anterior follicle cells ^25^. Further it has been reported that the TGF beta signaling regulates the timing of this transition ^7^. The transitioning of cuboidal to squamous fate is presumed to be a two-step process where the lateral membrane of the cuboidal cells shrinks followed by the stretching of flattened follicle cells ^26^. The flattening of the cuboidal follicle cells is aided by the removal of lateral protein Fas2 by the Ser/Thr kinase Tao ^27^. Though we know that the removal of Fas 2 protein is an important step for the initiation of the shape change and subsequent growth of the AFCs to acquire squamous fate, it is not very clear what other factors may be regulating this step.

In this study, we report a novel role for growth-promoting TOR signaling in mediating the shape change of cuboidal follicle cells to squamous cell fate. Employing fly genetics, live cell imaging, and immunohistochemistry approaches, we demonstrate that TOR complex 1 affects the shape transition of anterior cuboidal follicle cells to squamous fate. Our data suggest that TOR functions through the Serine threonine kinase protein Par-1 to mediate endocytosis in AFCs that are undergoing shape change. We report that Par-1 negatively affects the internalization of Fas2 in cuboidal follicle cells thus delaying their shape transition to squamous fate. Interestingly, our results uncover a novel aspect of the TOR function that is probably independent of its growth-promoting activity but affecting the temporal aspect of cell shape change. Altogether the results will further our understanding of how epithelial cells changes shape and will have broad implications in understanding both development and diseases in the metazoans.

## RESULTS

### Cuboidal-To-Squamous Cell Transition Involves a Dramatic Increase in Cellular Size and Exhibits Enriched TORC1 Reporter Activity

Previous studies have shown that the stage 8 cuboidal follicle cells undergo a conspicuous increase in cell size to acquire flattened squamous cell fate in the developing egg chambers ^7, 21, 25–27^. Consistent with the previous report, we also observed that the area of squamous cells is approximately 30 times (1316±87.1 SEM µm^2^) than that of their cuboidal precursors (26.49±1.915 SEM µm^2^) (Fig 1A). Although the area comparison between these two cell types can serve as a proxy for judging changes in cell size, we reasoned that simple redistribution of cuboidal cell volume can also generate a squamous cell fate. This can be facilitated by a substantial decrease in lateral height coupled with a concomitant increase in the surface area of the cuboidal cells. To verify this possibility, we calculated the volume of the cuboidal and squamous cells by multiplying their surface area by their respective height. We observed that the squamous cells have a larger volume (7147± 661.9 SEM µm^3^) compared to the cuboidal follicle cells (244.7±18.01 SEM µm^3^) (Fig 1B). Since the squamous cells are 30 times larger than their precursors, it suggested that the shape transition of AFCs in the late stage 8 egg chambers is associated with a significant increase in their cell size. Given that TOR has been reported to control the growth of follicle cells in developing egg chambers, we were curious to examine whether the growth-promoting TOR kinase plays a role in mediating the shape transition of the AFCs ^28^. First, we examined the status of TOR signaling in the developing egg chambers. To check this, we stained the egg chambers with antibody to evaluate the phosphorylation status of the Ribosomal protein S6 (pS6) that has been routinely used as a readout for TOR activity ^29, 30^. Unlike the late-stage egg chambers where the majority of follicle cells exhibit TOR activity, we observed a patchy expression of pS6 in the follicle cells of early-stage egg chambers (Fig 1C-E). A similar mosaic pattern of pS6 distribution has been reported in the *Drosophila* wing discs as well ^30^. Incidentally, we also observed that the stage 9 egg chambers exhibit spatial enrichment of pS6 staining in the anterior and posterior follicle cells (Fig 1F-H). As the anterior expression domain of pS6 coincides with the cuboidal cells that undergo shape transition to squamous fate, we wanted to check if the observed TOR activity in the AFCs has any role in mediating shape change of follicle cells in the developing egg chambers. First, we employed an RNAi approach to down-regulate TOR function to evaluate its role in the AFCs. Using FLIPOUT mediated system, we randomly generated mosaics of TOR^RNAi^ overexpressing clusters in the follicle cells. The cuboidal-to-squamous-shape transition of AFCs is a two-step process involving the shrinkage of the lateral membrane (flattening) followed by stretching of the flattened cells to acquire the squamous fate ^26^. The TOR^RNAi^ clones were identified by GFP expression and unlike the control where the AFCs have completely stretched, TOR-depleted AFCs were smaller and exhibited a defect in stretching in the stage 10 egg chambers (Control: 768.3±38.01 SEM µm^2^; TOR^RNAi^: 374.4±29.9 SEM µm^2^) (Fig 1I-O). In addition, we observed that the internuclear distance between the AFCs nuclei was much lower than that of the controls, (Control: 30.91±1.5 SEM μm, n=58; TOR^RNAi^: 20.64±1.6 SEM μm, n=58) suggesting that the stretching process during the shape transition was significantly impeded (Fig S1A). We also observed that follicle cells overexpressing TOR^RNAi^ exhibited lower levels of pS6 staining, suggesting that the TOR^RNAi^ construct is indeed depleting the TOR function (Fig S1B-D). Altogether the result above suggested that TOR plays a role in mediating the shape change of the AFCs.

**Figure 1:**
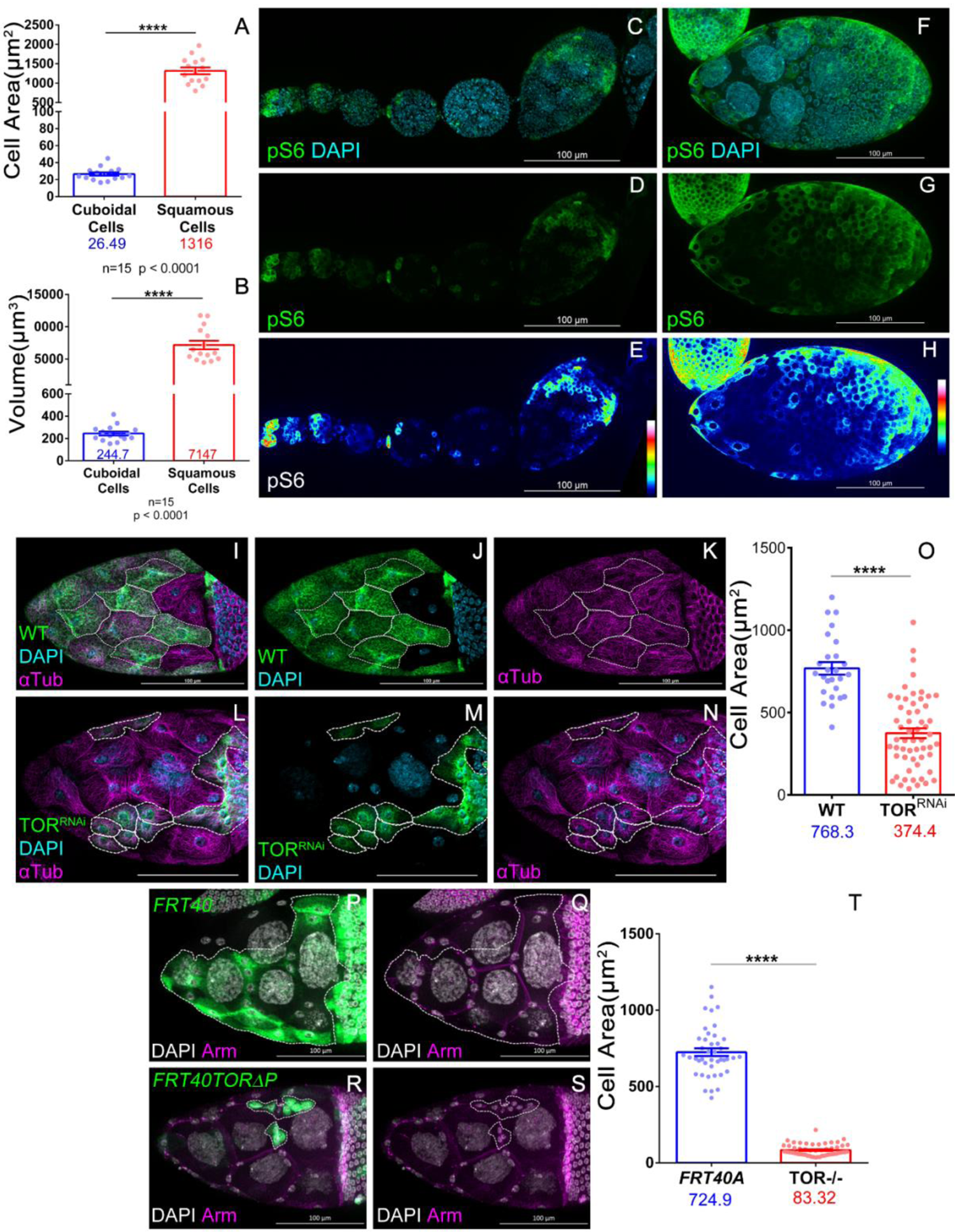
Requirement of TOR during Squamous Cell Morphogenesis. **(A)** Area Comparison of Cuboidal Cells and Squamous Cells. **(B)** Volume Comparison of Cuboidal Cells and Squamous Cells. **(C-E)** Distribution of pS6, TORC1 activity reporter in early oogenesis (Germline to S7/8). Note the patchy distribution of pS6 throughout oogenesis. **(F-H)** Distribution of pS6 in Stage 9 egg chamber. pS6 is enriched in the anterior cells transitioning into squamous cells. **(I-O)** Comparison of Cell Area in Control **(I-K)** and TOR^RNAi^ **(L-M)** expressing anterior follicle cells marked by Flipout Gal4 mediated mCD8GFP expression. **(O)** Plot comparing the cell area in control flipout cells and TOR^RNAi^ expressing Flipout cells. Mean values indicated below each genotype in mm^2^. **(P-T)** Comparison of cell area in Control **(P-Q)** and *TOR^ΔP^* **(R-S)** mutant anterior follicle cell clones, marked by mCD8GFP expression. **(T)** Plot comparing the cell area in control and *TOR^ΔP^* mutant clones, average values indicated below genotypes. Error bars indicate SEM. **\*\*\*\*** indicates p-value < 0.0001 in Students’ t-test with Welch’s Correction

To validate the RNAi phenotype, we employed the Mosaic Analysis with a Repressible Cell Marker (MARCM) to generate homozygous *TOR^ΔP^* mutant ^31, 32^ clones in the follicle cells. The purpose was to check whether the *clones of the TOR^ΔP^* mutant (null allele) clones impede the shape change associated with the AFCs. Consistent with our expectation, we observed *TOR^ΔP^* mutant AFCs were smaller in size compared to the controls phenocopying the TOR ^RNAi^-induced phenotype. The average area of the *TOR^ΔP^* mutants was 83.32±5.0 SEM μm^2^ (n=55) compared to 724.9±25.4 SEM μm^2^ (n=55) observed for the controls (Fig 1P-T). As TOR-depleted follicle cells exhibit reduced cell size, our data suggest that TOR plays a role in mediating cuboidal to squamous shape change of AFCs during *Drosophila* oogenesis. As a subset of anterior follicle cells undergoes active movement during egg chamber development, we were curious to check if the shape-change defect exhibited by TOR depletion was not an outcome of altered follicle cell fate specification.

### TOR depletion does not affect follicle cell proliferation or their fate

During *Drosophila* oogenesis, the shape change of AFCs coincides with the previtellogenic to vitellogenic transition of developing egg chambers. As the previtellogenic phase is associated with follicle cell proliferation, we wondered if the stretching defect observed in the TOR-depleted follicle cells was due to the excessive proliferation of the follicle cells itself. Unlike the vitellogenic phase, the follicle cells of the previtellogenic egg chambers retain phospho histone 3 (PH3) as they are undergoing active proliferation. Satisfyingly we observed the absence of PH3 marker in the TOR-depleted follicle cells (*GR1*-GAL4 / *UAS TOR^RNAi^*) till Stage 6 egg chambers suggesting that they have already ceased mitosis (Control- 7.8±0.85 SEM; TOR^RNAi^-6.7±0.47 SEM) (Fig 2A-G). However, we did observe 1 or 2 puncta of PH3 in the posterior follicle cells of Stage 7 egg chambers (33%, n= 18) (Fig S2A). This suggests that shape transition defects associated with TOR-depleted anterior follicle cells may not be a direct outcome of excessive proliferation of the cuboidal epithelial cells. Given that the TOR depletion phenotype was not arising due to excessive proliferation of AFCs. Next, we evaluated the fate of the AFCs exhibiting shape change defect.

**Figure 2:**
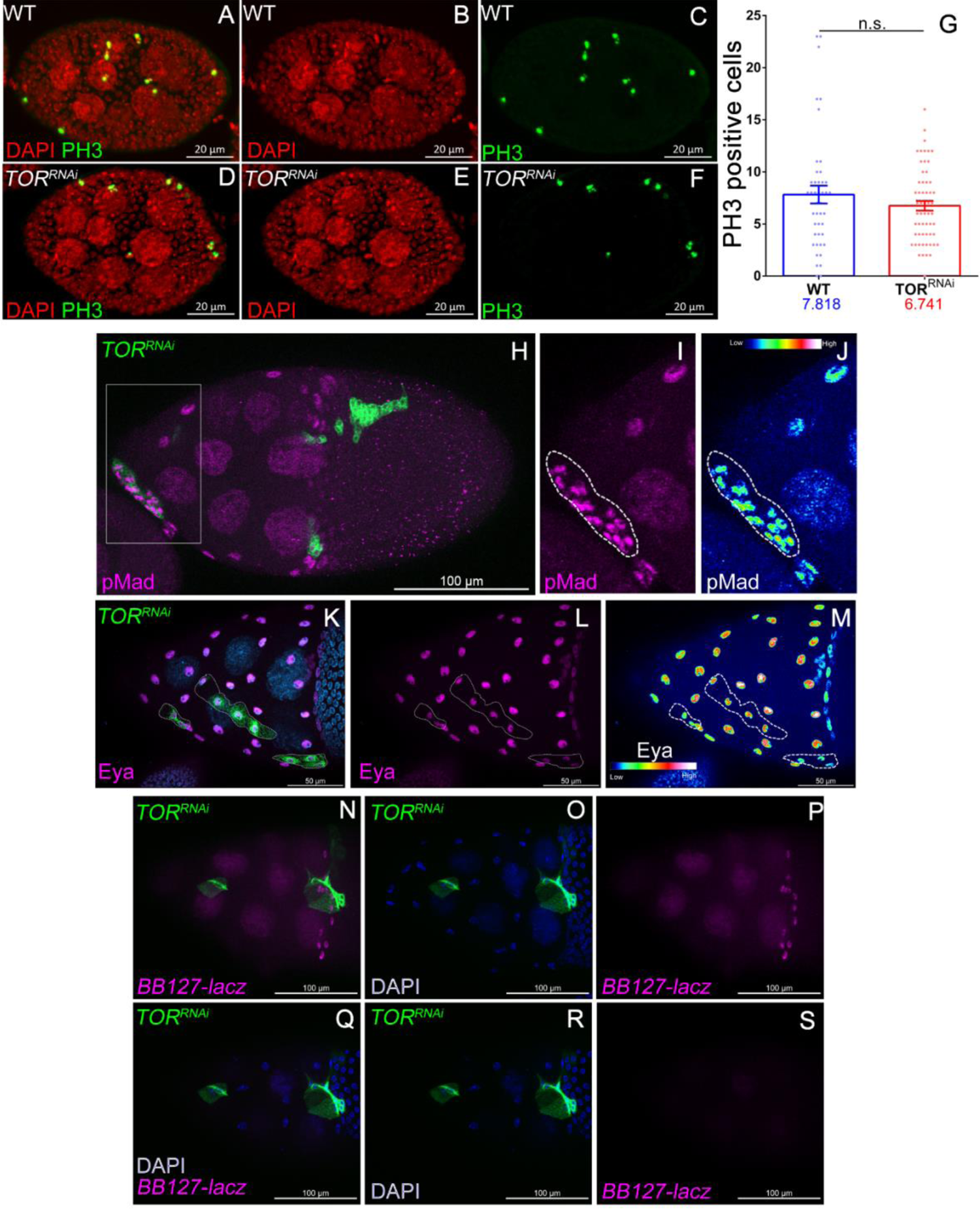
TOR depletion in anterior cells does not alter the cell fate. **(A-G)** Comparison of PH3 staining in *GR1*-Gal4 driven Control and TOR^RNAi^ follicular epithelium. PH3 count across S4-S6 was unaffected between Control and TOR depleted follicle cells. **(H-J)** TOR depleted follicle cells are positive for pMad staining. **(K-M)** TOR depleted follicle cells are positive for Eya staining. **(N-S)** TOR depleted follicle cells do not carry the centripetal marker *BB127-LacZ*. **(N-P)** is a z-projection of planes including anterior follicle cells and centripetal cells marked by *BB127-LacZ* expression (magenta). **(Q-S)** Same egg chamber with shorter z-projection showing absence of *BB127-LacZ* marker in TOR depleted follicle cells marked by mCD8GFP expression. Error bars indicate SEM. ns = not significant, in Students’ t-test with Welch’s correction.

We know that Dpp signaling regulates the timing of the shape transition of the cuboidal AFCs to squamous fate ^7^. So, we were curious if the phenotype associated with TOR depletion follicle cells is not just an outcome of AFCs stalled in early oogenesis due to lack of Dpp signaling. As Dpp activation leads to phosphorylation of Mad transcription factor (pMAD), we observed that the TOR-depleted AFCs do stain for pMAD like the control follicle cells (Fig 2H-J) suggesting that the shape change defect associated with TOR depletion is not an outcome of follicle cells being deprived of Dpp signaling^33^.

Next, we evaluated the fate of the TOR-depleted follicle cells *per se* that exhibit the shape change defect. This was because we know that apart from the cuboidal to squamous cell shape change, there are two significant shape transition events associated with remaining anterior cuboidal cells. The cuboidal cells over the nurse cells slightly posterior to AFC migrate towards the oocyte ^21^. One set of cells called the centripetal cells, laterally elongate and move inward between the nurse cells and oocyte. Another set of cells move posteriorly and elongate to columnar fate commonly referred to as main body follicle cells (MBFC). To check the possibility that the TOR-depleted cells that exhibited shape change defects were not the unmigrated MBFCs or the centripetal cells, we evaluated the status of molecular markers that label these distinct follicle cell populations. Eya is a transcription factor, that marks all follicle cells including cuboidal cells till stage 8 and squamous and centripetal cells in stage 10A egg chambers ^34^. We observed that TOR-depleted follicle cells that exhibit smaller cell size and shape change defects continue to express the Eya protein, suggesting that the overall fate of follicle cells is not impeded nor these cells are unmigrated MBFCs (Fig 2K-M). To further check if these TOR-depleted follicle are the unmigrated centripetal cells, we examined the status of another marker *BB127-Lacz*. The expression of *BB127-Lacz* categorically marks the nurse cells and the centripetal cells^35^. Satisfyingly we didn’t observe any expression of *BB127-Lacz* in the AFCs exhibiting shape change defects suggesting that the TOR-depleted AFCs are neither the centripetal cells (Fig 2N-S). Altogether the results above suggest that the phenotype associated with TOR depletion is indeed an outcome of the shape change defect of AFCs and not a consequence of unmigrated centripetal cells or the MBFCs.

As remodelling of adherens junction is critical for facilitating shape change of AFCs, we were curious to check if the shape change defect was associated with the inability of TOR depleted AFC to modify their adherens junctions.

### TOR-depleted AFCs exhibit delay in stretching in real-time analysis

Since cell shape change is a dynamic process, we employed live cell imaging to systematically examine how TOR signalling is affecting the shape change of anterior follicle cells. First, we chose the *GR1*-GAL4 driver for overexpressing TOR^RNAi^ as it gave modest defect in the area of shape transition AFCs compared to the control cells (Control cells had an area of 863.6±27.6 SEM µm^2^, while TOR^RNAi^ cells had 455.5±22.2 SEM µm^2^ (Fig S3A-C). In time-lapse imaging, we particularly focussed on AFCs of stage 8-9 egg chambers and used time-lapse imaging to capture the transformation to a squamous shape. We labelled the lateral domain of follicle cells with the endogenous DE-Cadherin:GFP ^36^. As the egg chamber development progressed, the AFCs exhibited a dramatic increase in their surface areas with a concomitant decrease in the intensity of the DE-Cadherin: GFP. This decrease in DE-Cadherin:GFP signal may suggest that the lateral membrane is getting shortened as the AFCs are undergoing shape change. Unlike the control AFCs, the intensity of the DE-Cadherin:GFP perdured for a much longer duration in the TOR-depleted AFCs suggesting retention of a significant portion of their lateral membrane (Fig 3A^1^-B^6^, Movie 1-2).

**Figure 3:**
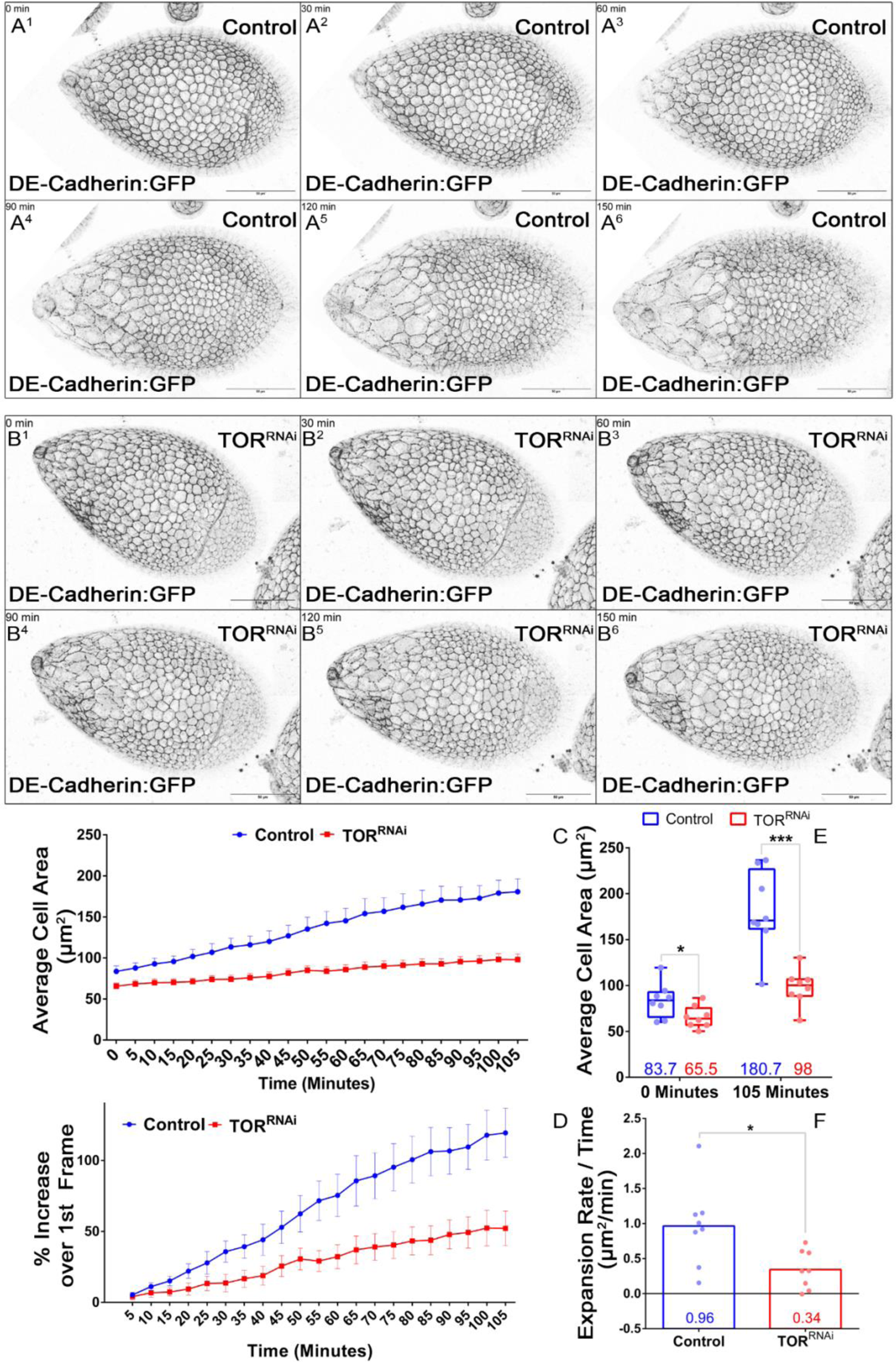
Live imaging reveals slower expansion rate of anterior follicle cells upon *GR1*-Gal4 mediated TOR Depletion. **(A^1-6^)** Montage of control egg chambers of genotype DE-cadherin:GFP; *GR1*-Gal4/+; Note the anterior follicle cells expanding into the squamous cells. Represented 30minutes apart. Scale bars denote 50 μm. **(B^1-6^)** Montage of TOR depleted egg chambers carrying DE-cadherin:GFP; *GR1*-Gal4/ TOR^RNAi^. Note, anterior follicle cells exhibiting incomplete and non-uniform expansion upon TOR depletion. Represented 30 minutes apart. Scale bars denote 50 μm. **(C)** Comparison of average cell area in each egg chamber at each time point (5 min apart) reveals decreased expansion of the anterior cell population. Error bars indicate SEM. **(D)** Comparison of % increase in area of anterior squamous cells of consecutive time points with respect to first time point. **(E)** Comparison of average anterior cell area across Control and TOR^RNAi^ genotype, at initial time point and at the end of 105 minutes, reveals defect in expansion of anterior follicle cells upon TOR depletion. Plot represented with box and whiskers, mean with min to max (all points). **(F)** Comparison of average of expansion rate across each egg chamber in control and TOR depleted condition. Error bars indicate SEM. **\*** Denotes p<0.05 in Students’ t-test with Welch’s correction.

Next, we sequentially mapped the outline of the expanding AFCs by following the DE-Cadherin:GFP and derived the surface area frame by frame in the developing egg chamber. Stage 5-9 egg chambers are known to exhibit rotation around their anterior-posterior axis ^37^. Consequently, any area obtained for cells during the rotation phase will be influenced by egg chamber movement. Hence, we limited our area analysis to time frames, where rotation had visually stopped.

We first plotted the mean area of expanding AFCs in control and TOR-depleted background, over 105 minutes (Fig 3C). Control cells started with an average area of 83.7±6.7 SEM µm^2^ compared to 65.5±4.2 SEM µm^2^ under TOR depletion. In the end, control cells displayed an area of 180.7±15.6 SEM µm^2^ compared to 98±6.9 SEM µm^2^ observed in the TOR depletion condition (Fig 3E). These numbers correspond to an average increase of 2.2-fold in the area of control follicle cells, compared to an increase of 1.5-fold observed in TOR-depleted follicle cells (Fig 3C,E). Second, we plotted the % expansion of AFCs in each frame in reference to the first frame. Control AFCs exhibited around 119.5±17.3 SEM % increase in area over the first frame, compared to a 52.1±12.1 SEM % increase observed in the TOR-depleted follicle cells (Fig 3D). Finally, we measured the average rate of area expansion of AFCs, to the first time point. Control cells exhibited an average expansion rate of 0.96 µm^2^/min compared to 0.34 µm^2^/min under TOR depletion (Fig 3F). In sum, all the data above suggests that TOR depletion leads to defective and slower expansion of anterior follicle cells into squamous type.

Altogether, the real-time analysis suggests that TOR-depleted AFCs are deficient in expanding their surface and retain their lateral membrane for a longer duration than the control. This prompted us to compare the status of lateral markers between the wildtype (WT) and TOR-depleted AFCs.

### TOR Regulates Squamous Cell Morphogenesis by affecting Fas2 downregulation

A critical step during the morphogenesis of cuboidal cells to squamous fate involves the timely removal of lateral cell adhesion molecule, Fasciclin 2. Fasciclin 2, the *Drosophila* homolog of Neural-Cell Adhesion Molecule (N-CAM), is endocytosed out from the lateral membrane of the AFCs in stages 7-8 egg chambers to facilitate their shape change ^27, 38^ (Fig 4A-B). By the end of stage 10, Fas2 is undetectable in the egg chambers except at the interface between the pair of polar cells (Fig 4C-D). Since pS6 is detected around the same time when Fas2 removal is initiated in the anterior follicle cells, we were curious to test the possibility if TOR activity is essential for promoting the endocytic removal of Fas2 and thereby facilitating the transition of cuboidal cells to squamous fate. We reasoned that if TOR is affecting cuboidal-to-squamous cell shape transition by regulating Fas2 removal, then TOR depletion would exhibit retention of Fas2 in the AFCs (Fig 4E-E’’). We indeed observed that TOR-depleted anterior follicle cells exhibit delayed Fas2 removal compared to the nearby control follicle cells. This was also recapitulated when *TOR^ΔP^* mutant follicle cells were generated by the MARCM technique suggesting that TOR depletion indeed impedes Fas2 removal from the lateral membrane of follicle cells (Fig 4F-F’’). We also knocked down TOR function in the background of endogenously tagged Fas2^39739^ and observed a similar higher amount of Fas2:GFP in the clonal cells compared to neighboring WT cells (Fig S4A-C). To further check if the delay in Fas2 downregulation was indeed the cause of the morphogenesis defect associated with AFCs, we attempted to rescue the defective cell shape change phenotype by reducing the levels of Fas2 in TOR depleted background. When we introduced the amorphic allele, *fas2^EB112^*,/+ in TOR mutant follicles cells, we could partially rescue the cell size of TOR-depleted AFCs undergoing shape change (*TOR^ΔP^*: 69.82±5.4 SEM µm^2^, +/ *fas2^EB112^; TOR^ΔP^:* 182.1±13.6 SEM µm^2^) (Fig 4G-I) ^40^. Since Fas2 reduction can rescue the cell size of TOR depleted AFCs, it suggested that TOR mediates Fas2 down-regulation in the cuboidal AFCs as they transition their shape to the squamous fate.

**Figure 4:**
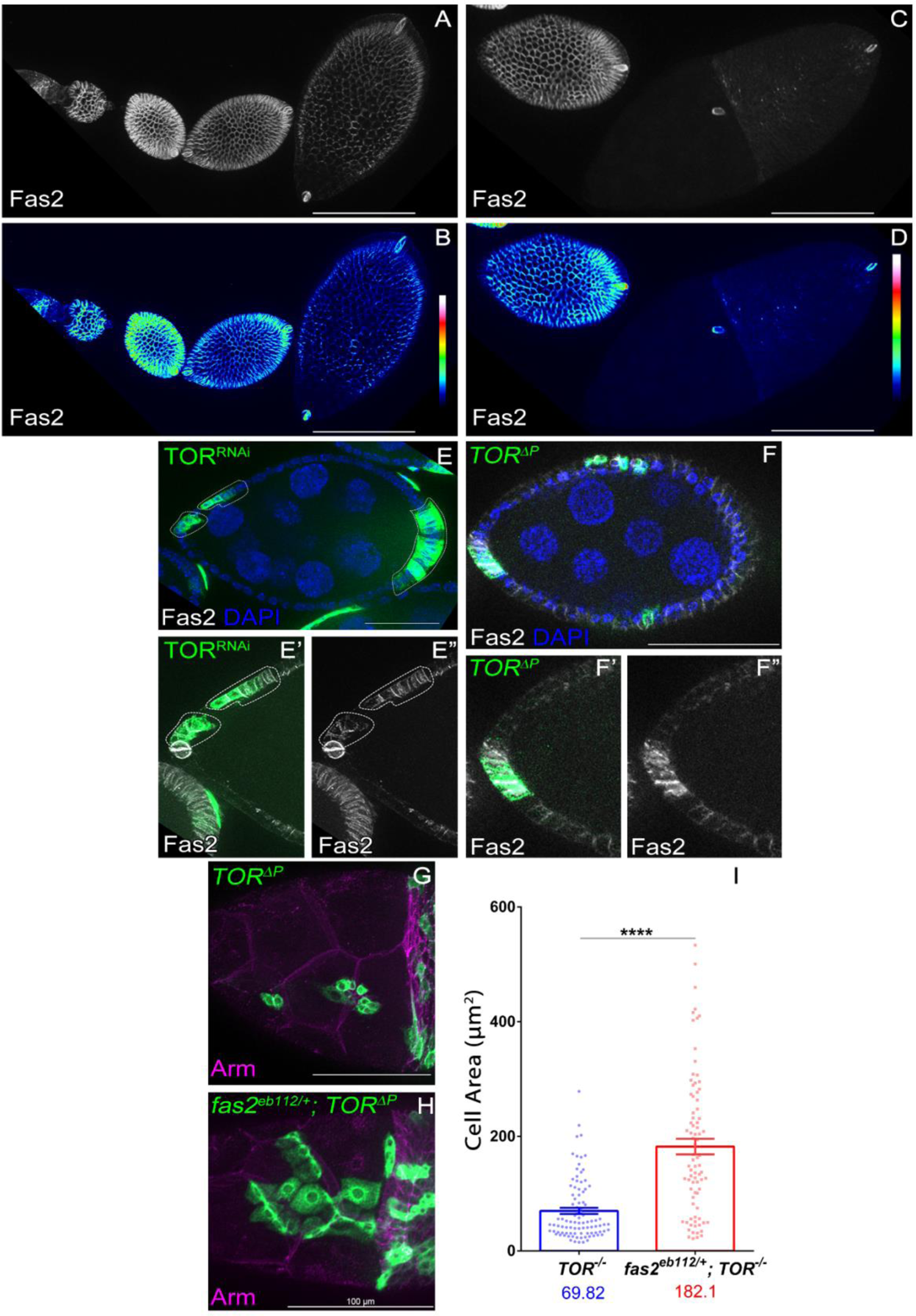
Improper Fas2 depletion is the cause of morphogenesis defects observed upon TOR Depletion. **(A-B)** Distribution of Fas2 in early – Stage 7 egg chambers. Distribution of Fas2 in **(A)** White and **(B)** as heatmap. Note presence of Fas2 across the lateral side of all follicle cells. **(C-D)** Comparison of Fas2 in Stage 6/7 egg chamber (left) and Stage 10 egg chamber (right). Fas2 represented in **(C)** as White and **(D)** as Heatmap. **(C)** Note the presence of Fas2 in the smaller Stage 6/7 egg chambers (on left). **(D)** Negligible Fas2 expression at the anterior end of the larger Stage 10 egg chamber (on right). The posterior cells display some degree of Fas2. The polar cells pairs continue to display Fas2 – one at oocyte border (as part of border cells) and other at the posterior end of the egg chamber. **(E-E’’’)** TOR^RNAi^ expressing anterior cells display delayed Fas2 (white) removal. All clonal population is marked by GFP expression (green). **(F-F’’)** *TOR^ΔP^* mutant clones (green) exhibit similar delayed Fas2 (white) removal phenotype. **(G-I)** Genetic removal of Fas2 (*fas2^eb112/+^* allele) in *TOR^ΔP^* rescues the cell area defects observed upon *TOR^ΔP^.* **(I)** Comparison and Plot of cell area across *TOR^ΔP^* and *fas2^eb112/+^; TOR^ΔP^.* Error bars indicate SEM. **\*\*\*\*** denotes p-value < 0.0001 in Students’ t-test with Welch’s correction.

### TOR mediates endocytosis in the follicle cells

Previous studies have revealed that endocytosis is one of the downstream targets of the TOR signalling pathway ^41, 42^. Given that internalized Fas2 has been shown to colocalize with Rab5 in the AFCs, we first assessed the status of endocytosis in the TOR-depleted follicle cells ^27^. To check this, egg chambers carrying *TOR^RNAi^* clones were intermittently exposed to tracer dye TR-Avidin and analyzed for dye internalization under live conditions. Under epifluorescence/Apotome based microscopy, we observed a dramatic localization of TR-avidin at the interface between posterior columnar cells and the oocyte, while in TOR-depleted follicle cells internalized dye was reduced. This suggests that TOR-depleted follicle cells exhibit a defect in endocytosis (Relative levels – 1.00 in control and 0.68±0.03 SEM in *TOR^RNAi^*, n=31 egg chambers) (Fig S5A-D) Given that TOR signaling affects endocytosis in the follicle cells, we were curious to evaluate if the TOR and endocytosis function together to modulate the shape change of AFCs. Hsc70-4 is a clathrin uncoating ATPase that has been previously shown to genetically interact with TOR in the eye imaginal discs ^42, 43^. So, we analyzed the shape change defect in FLIP OUT overexpression clones of *TOR^RNAi^* in wild type and *Hsc70-4 ^L^*^3929^ heterozygous null allele background. We compared cell size in *TOR^RNAi^* expressing clones in presence of endocytic regulator Hsc70-4/+ (*hsc70-4^L39^*^29^/+) mutant allele to clones expressing only TOR^RNAi^. The TOR-depleted follicle cells in the *hsc70-4^L3929^/+* background were smaller (248±26.03 SEM μm^2^) than that observed in TOR^RNAi^ (538±31.5 SEM μm^2^) or WT background (1138 ± 41.28 SEM µm^2^) (Fig 5A). Next, we checked if endocytosis *per se* can regulate this morphogenetic event. To assess this, we perturbed the function of the following endocytic regulators, Shibire (vesicle pinching) and Hsc70-4 (clathrin uncoating ATPase), by overexpressing their transgenes, Shi^DN^, Shi^TS^ and Hsc70-4^RNAi^, respectively in the AFCs. We observed a significant decrease in the follicle cell size in this background compared to the control follicle cell clones with the Hsc70-4^RNAi^ exhibiting the strongest phenotype among all the endocytotic regulators tested (Control-723.5±19.3 SEM µm^2^; Hsc70-40^RNAi^-152.9±15.3 SEM µm^2^; Shi^DN^-386.2±28.2 SEM µm^2^; Shi^TS-GFP^-340.3±17.7 SEM µm^2^) (Fig 5B-H). We also evaluated the status of Fas2 in the Hsc70-4^RNAi^ follicle cell clones and pleasingly we observed retention of Fas2 in the clones exhibiting shape change defect (Fig 5I-J). Thus, we conclude that endocytosis regulates the shape change of AFCs to squamous cell fate. Overall, our results above suggest that TOR signaling functions with endocytosis regulators to modulate squamous cell morphogenesis of the AFCs in the developing egg chambers. Next, we were curious to find out how TOR was modulating endocytosis to affect the shape transition of anterior follicle cells.

**Figure 5:**
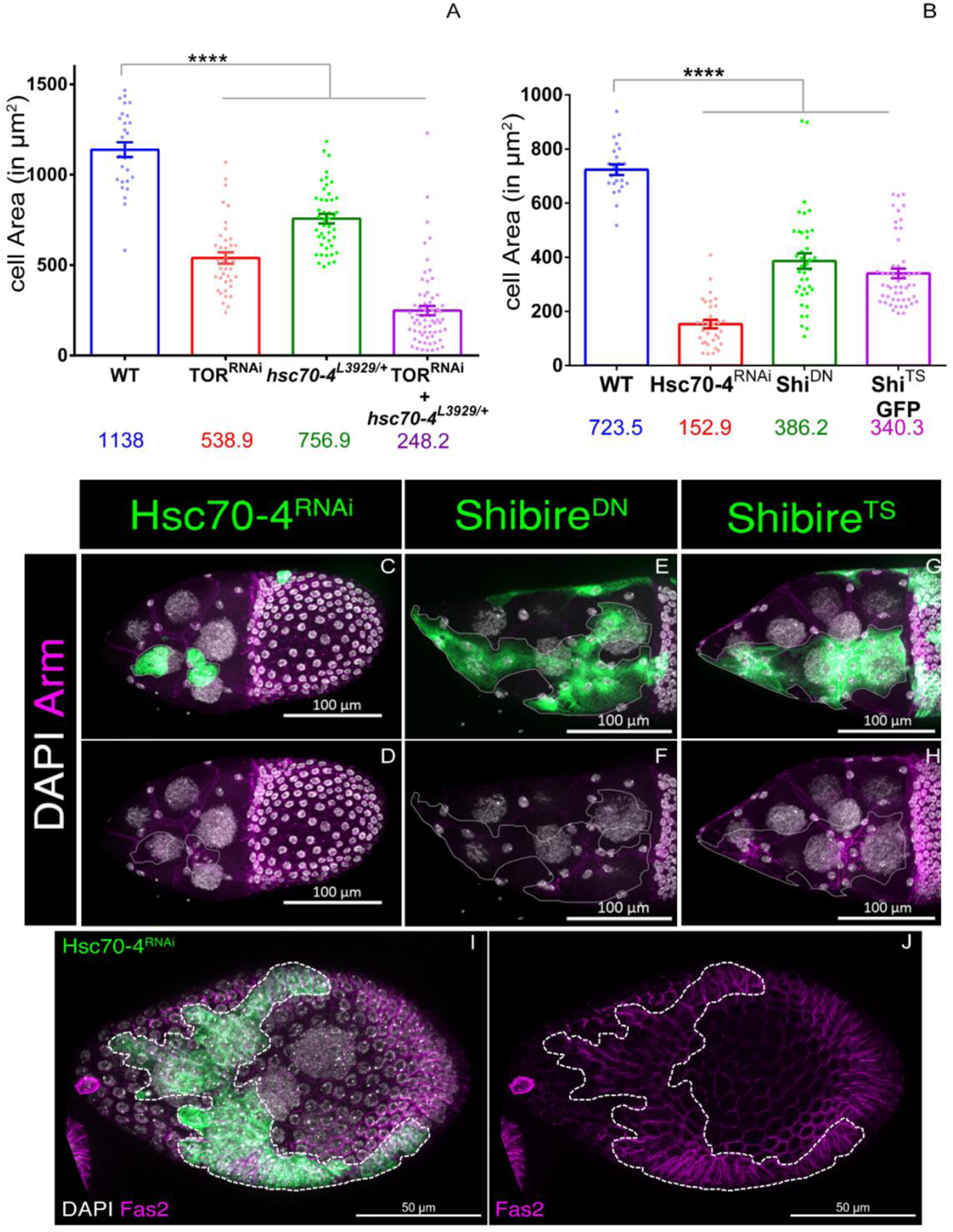
Endocytosis and TOR mediates cuboidal-to-squamous transition. **(A)** Genetic Removal of Hsc70-4 increases the cell size defects observed upon TOR depletion. Cell area comparison in genotypes - *WT*, *TOR^RNAi^*, *hsc70-4^L3929/+^*, *TOR^RNAi^* + *hsc70-4^L3929/+^*. **(B-J)** Perturbation of endocytosis alone affects proper cuboidal to squamous cell shape transition. **(C-D)** Anterior follicle cells expressing Hsc70-4^RNAi^ construct, marked by mCD8GFP (green). **(E-F)** Anterior follicle cells expressing Shibire^DN^ (dominant negative) construct, marked by mCD8GFP expression (green). **(G-H)** Anterior follicle cells expressing Shibire^TS^ (temperature sensitive) construct, marked by mCD8GFP expression (green). **(B)** Plot representing the cell area in the mentioned genotypes. **(I-J)** Timely Fas2 (magenta) removal is affected in follicle cells expressing Hsc70-4^RNAi^, marked by mCD8GFP expression (green). Error bars denote SEM. **** represent p-value < 0.001 in Students’ t-test with Welch’s correction.

### TOR complex1 affects the shape transition of AFC like the TOR mutant

TOR kinase functions through two distinct complexes TOR Complex 1 and 2 (TORC1 and TORC2) to affect diverse cellular processes linked to cell growth and autophagy ^8,9,44^. To further understand how TOR is affecting anterior follicle cell morphogenesis, we evaluated which of the TOR complexes was mediating the shape change associated with AFCs. The functional presence of the Regulatory Associated Protein of TOR (RAPTOR) marks activity for complex 1 while the activity of Rapamycin Insensitive Companion of TOR (RICTOR) can be used as a functional readout for complex 2 ^10, 11, 45, 46^. To check which of these complexes is indeed required to mediate the shape change of AFCs, we downregulated the function of RAPTOR and RICTOR genes independently in the follicle cells by RNA interference. Our premise was that the RNAi construct that impedes shape change of AFCs similar to that observed for the TOR mutants will indicate as to which TOR complex is involved in this process. First, we validated both the RNAi constructs themselves by evaluating pS6 levels in RAPTOR^RNAi^ overexpressing clones (Fig S6A-B) and levels of RICTOR mRNA in RICTOR^RNAi^ overexpression background (Fig S6K-L). In our experiment to delineate which TOR complex mediates the shape change of AFCs, we observed that RAPTOR^RNAi^ over-expressing clones resulted in diminished AFCs size resembling the TOR mutant phenotype. Unlike the control follicle cells (891.1±36.4 SEM µm^2^), the RAPTOR^RNAi^ overexpressing follicle cells were smaller (367.0±19.8 SEM µm^2^) and similar to TOR^RNAi^ (233.5±20.2 SEM µm^2^) (Fig 6A-C, F). However, follicle cells depleted of RICTOR function exhibited normal shape transition just like the control cells (784.6±30.2 SEM µm^2^) (Fig 6E,F). To further check if the shape change defect observed in RAPTOR depleted follicle cells is phenocopied at the molecular level. We examined the levels of Fas2 and consistent with our expectation we observed Fas2 retention in RAPTOR^RNAi^ clones, supporting our claim that reduced TORC1 activity is associated with the retention of Fas2 in the AFCs (Fig 6G-G’’). We didn’t observe any Fas2 retention in RICTOR-depleted AFC clones (Fig 6H-H’’). Unlike the RAPTOR^RNAi^ clones, down regulation of S6 kinase function by RNAi didn’t impede the shape change of the AFCs (1035±26.4 SEM µm^2^) (Fig 6D,F). In addition, overexpression of constitutively active S6 Kinase (267.2±16.4 SEM µm^2^) failed to rescue the TOR ^RNAi^ (330.4±18.5SEM µm^2^) induced shape change defect in the AFC (Fig S6J). Both the S6K^RNAi^ and S6K^CA^ constructs were validated by evaluating pS6 levels in the follicle cells (Fig S6C-I).

**Figure 6:**
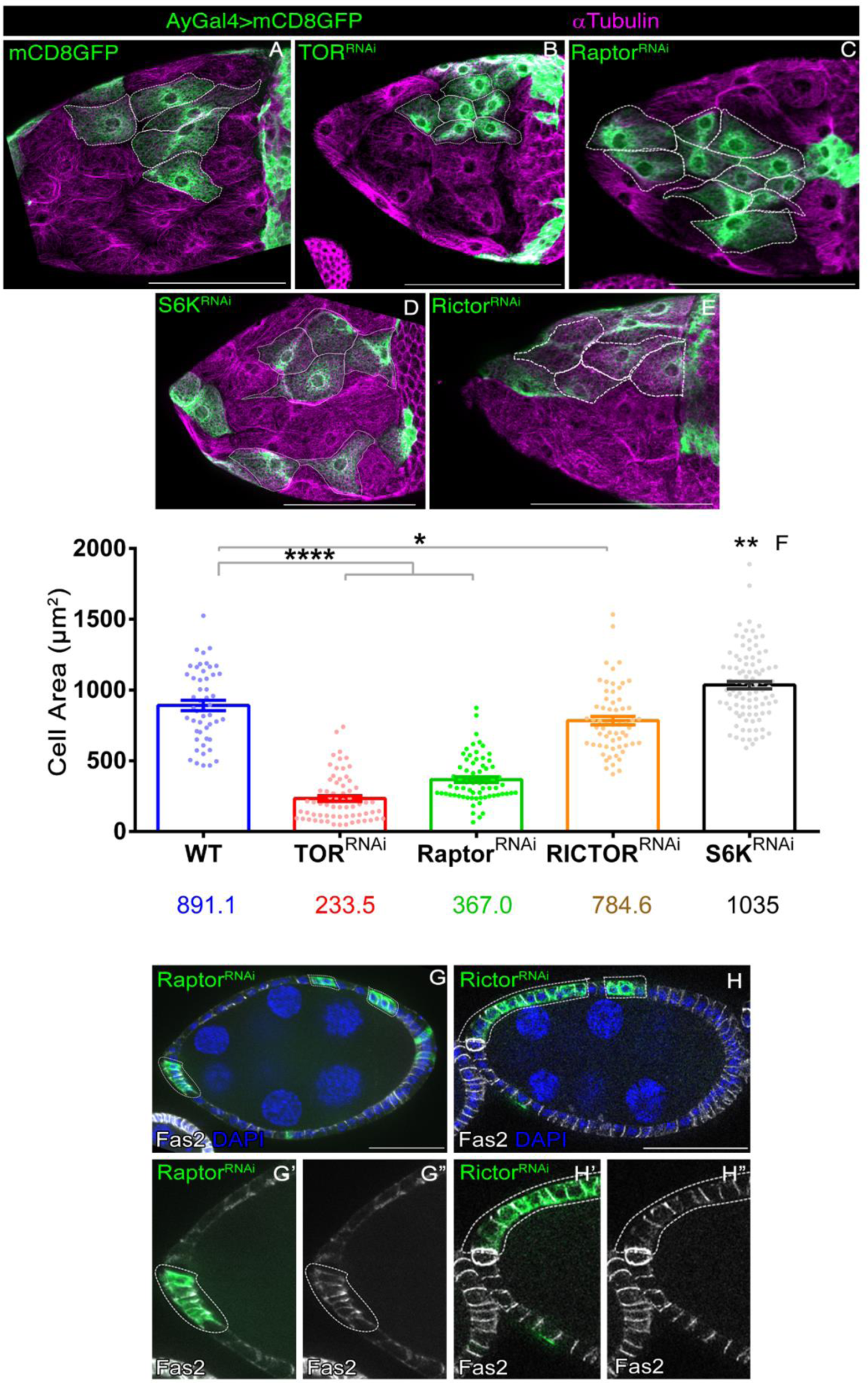
TORC1 primarily drives cell shape change in Drosophila follicular epithelium. **(A-E)** TOR Complex 1 primarily drives the cuboidal to squamous cell morphogenesis. Comparison of cell size in clonal populations (marked by mCD8GFP, green) – **(A)** mCD8GFP, **(B)** TOR^RNAi^, **(C)** Raptor^RNAi^, **(D)** S6K^RNAi^ **(E)** Rictor^RNAi^. **(F)** Plot comparison the cell size across the indicated genotypes. **(G-H’’’)** Raptor Depletion, but not Rictor Depletion affects timely Fas2 (white) removal from the lateral side. Clonal population is marked by mCD8GFP expression (green). **(G-G’’’)** Raptor^RNAi^ expressing anterior cells display delayed Fas2 removal. **(H-H’’’)** Rictor^RNAi^ expressing anterior cells does not exhibit delayed Fas2 removal phenotype. Error bars represent SEM. *, ** and **** represent p-value < 0.05, 0.01 and 0.0001 respectively in Students’ t-test with Welch’s correction

Altogether, the results above suggest that TOR complex1 (TORC1) mediates the shape change of the AFC during proper squamous cell morphogenesis and intriguingly it may function independently of S6 kinase. Next, we were curious to check how TOR complex 1 was mediating the shape change of AFCs during squamous cell morphogenesis.

### TOR impinges onto the endocytic pathway independent of REPTOR transcriptional regulation

The effect of active TOR signalling can be mediated through either transcriptional, translational, or post-translational regulation. One mechanism by which TOR mediates its role in cell growth is by modulating the activity of translation regulator S6 kinase. Interestingly in our experiments, any modulation of S6 kinase activity failed to exhibit any effect on the shape change of AFCs. Thus, we wanted to investigate if TORC1 was affecting the shape change of AFCs by modulating the transcription of downstream target genes. REPTOR is a transcription factor that mediates most of the altered transcriptional output observed upon TORC1 downregulation and is considered analogous to the FOXO transcription factor, a downstream target of AKT^47^. An active TOR kinase phosphorylates REPTOR and sequesters it in the cytosol ^47^. When TOR kinase is inactive, REPTOR is no longer phosphorylated, enters the nucleus, and generates an altered transcriptional profile, which has been shown to mimic similar TOR loss of function phenotypes ^47^. We overexpressed a constitutively active form of REPTOR (REPTOR^CA^) that cannot be phosphorylated by TOR and assessed for the morphogenesis defects of AFCs ^47^. We found that overexpression of REPTOR^CA^ (698±23.5 SEM µm^2^) does not completely phenocopy TOR loss-of-function phenotypes (Fig 7A-E). Nor does it exhibit any changes in the levels of Fas2 suggesting that TORC1 might be affecting the shape change of AFC independent of any transcriptional input (Fig 7F-H). This led us to ponder how TOR was modulating endocytosis of Fas2 in the AFCs.

**Figure 7:**
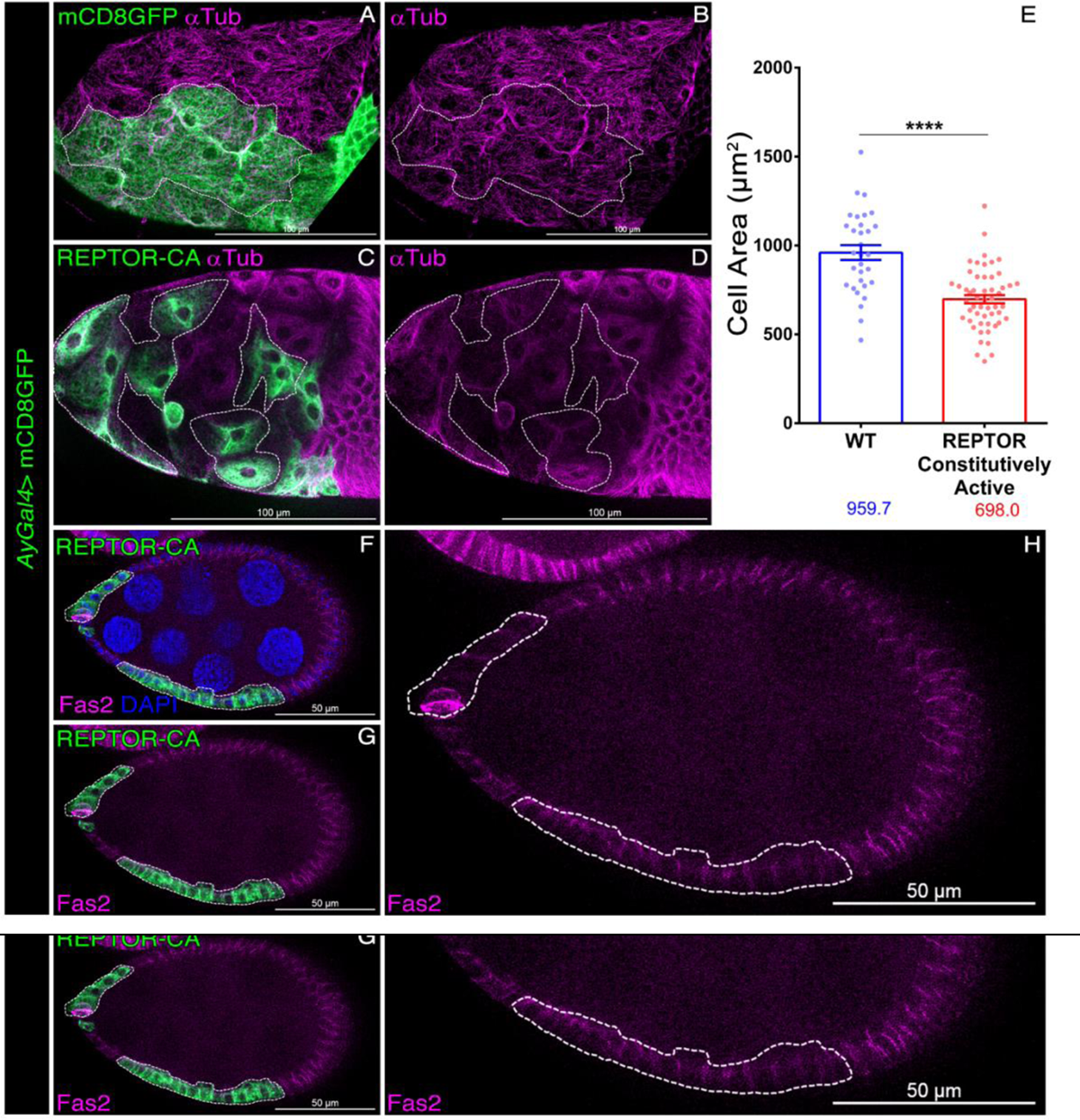
TOR regulates cell shape transition mostly independent of the transcription induction via REPTOR activity. **(A-E)** Induction of REPTOR Constitutively Active (CA) (green) construct does not majorly affect the size of anterior squamous cells. **(E)** Comparison of cell size in Control **(A-B)** and REPTOR-CA **(C-D)** expressing follicle cells. Error bars indicate SEM. **** denotes p-value in Students’ t-test with Welch’s correction. **(F-H)** REPTOR-CA expressing follicle cells, marked by mCD8GFP expression (green), does not exhibit Fas2 (magenta) retention phenotype.

### Par-1 genetically interacts with TOR to modulate Squamous Cell Morphogenesis

TOR has been shown to regulate endocytosis in a variety of organisms including yeast, flies, and rodents. In yeast, Nitrogen Permease Reactivator 1 (Npr1) is shown to regulate the cell surface display of amino acid importers via endocytic trafficking. Npr1, which is inhibited by TORC1 activity, is known to impede endocytosis, Since TORC1 promotes endocytosis in yeasts via inhibition of Npr1 ^48^. We decided to investigate whether a similar TOR-Npr1-Endocytosis axis exists in flies. To identify the fly homolog, we used the Npr1 protein sequence to perform a BLAST search in the fly proteome. Interestingly Par-1 polarity kinase was identified as the most significant hit in the BLAST searches (Fig S7A). However, this was unexpected as the KIN1/MARK family of proteins are the established Par-1 homologs in yeasts ^49, 50^. Interestingly, we observed that Par-1 (Par-1:GFP) and Fas2 exhibit overlapping expression along the lateral side of developing follicle cells in stage 8 chambers (Fig S7B-B”). Next, we checked how Par-1 affects endocytosis in the follicle cells by TR-avidin assay. Conspicuously, Par-1-depleted follicle cells accumulated higher TR-avidin compared to nearby control cells not expressing Par-1^RNAi^ suggesting that Par-1 negatively affects endocytosis in the developing follicle cells (Fig 8A-C). Having observed that Par-1 knockdown was promoting endocytosis, we next asked if the presence of Par-1 was linked to preventing premature removal of Fas2 from the lateral membrane of transitioning AFCs. To check this, we immuno-stained egg chambers expressing Par-1^RNAi^ with Fas2 antibody. We found a premature decrease in Fas2 during stages 6-8 of Par-1 depleted follicle cells, compared to the nearby WT cells (Fig 8D-G), suggesting that Par-1 negatively regulates endocytosis of Fas2 in the AFCs. This prompted us to test whether this antagonistic relationship between Par-1 and Fas2 can influence TOR-mediated regulation of cuboidal to squamous cell morphogenesis. To check this, we first examined the distribution of Par-1 and Fas2 on the lateral membrane of WT and TOR-depleted AFCs. The Par-1 and Fas2 exhibit diffuse staining in the follicle cell and line profile analysis demonstrated diffuse overlap among the two on the lateral membrane of follicle cells. Unlike the WT follicle cells, we observed a discrete membrane localized Par-1 with distinct enrichment at the apical and basolateral ends of the TOR-depleted follicle cells (Fig S7C-E). Since TOR-depleted follicle cells exhibit higher levels of Par-1 (Fig S7C-D’), we were curious if this was indeed the cause for the observed cell shape change defect observed in the AFCs. To test our hypothesis, we reduced Par-1 levels in TOR-depleted follicle cells and examined their size. Our premise was that if TOR was indeed negatively regulating Par-1 to aid the shape change of AFCs, then reducing Par-1 levels would rescue the size of the TOR-depleted follicle cells. Indeed, it was the case, as we observed larger cells in TOR-depleted AFCs in Par-1 heterozygous background than that observed in the TOR ^RNAi^ background alone (281 ±11.2 SEM µm^2^ in TOR^RNAi^ vs 456.7±12.9 SEM µm^2^ in *par-1^27C1^* /+; TOR^RNAi^) (Fig 8I). Our data above suggests that TOR negatively regulates Par-1 thus modulating the shape change of AFCs from cuboidal to squamous fate. Altogether, our results suggest that TORC1 functions through Par-1 to facilitate Fas2 removal in the AFCs that are undergoing shape change. Since TOR and Par-1 can modulate endocytosis, we believe that TOR may be assisting lateral membrane disassembly by restricting Par-1 to the lateral surface and facilitating Fas2 endocytosis.

**Figure 8:**
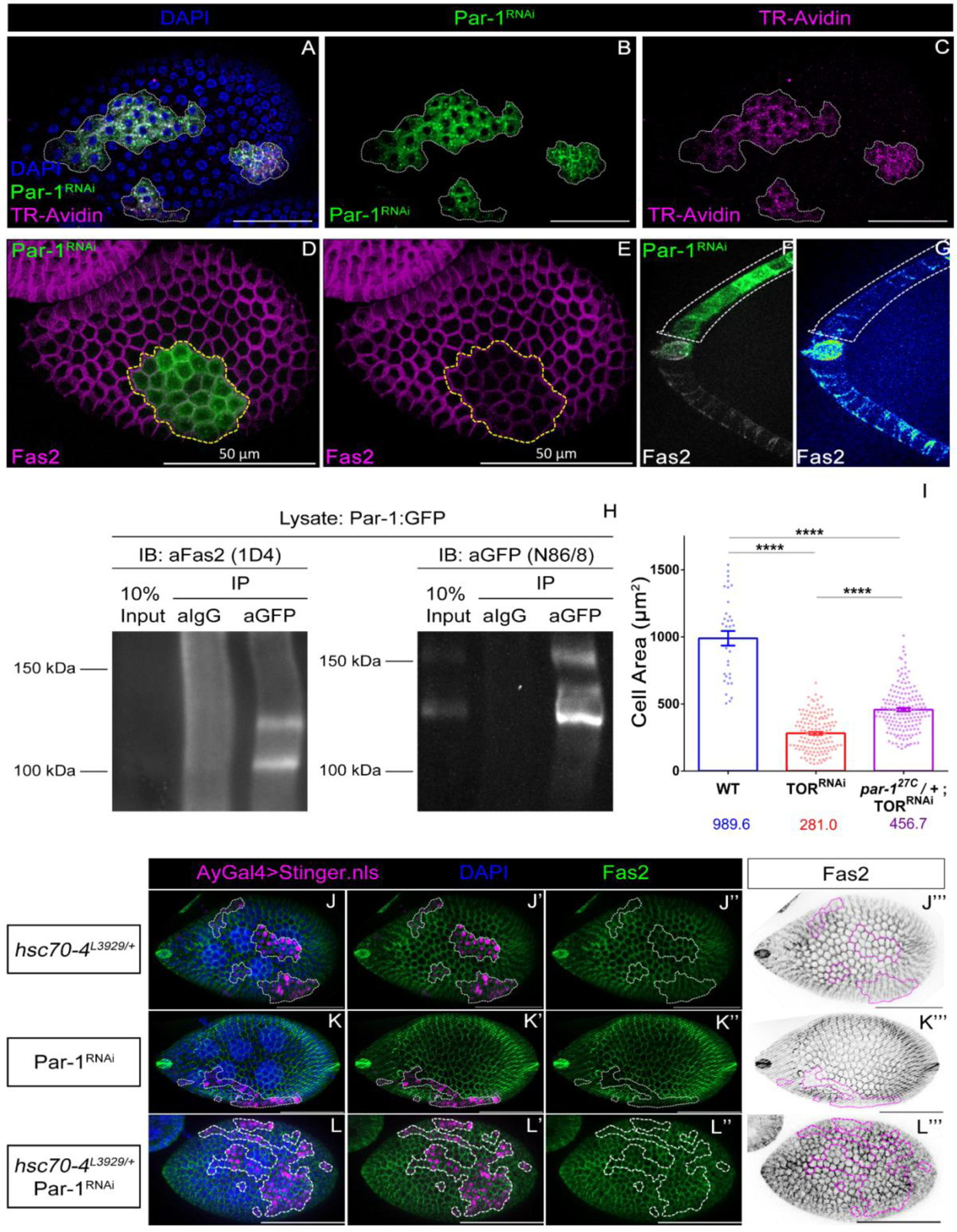
TOR regulate cuboidal to squamous cell transition by antagonising Par-1 mediated negative regulation of Endocytosis. **(A-C)** TR-avidin (magenta) uptake assay in follicle cells expressing Par-1^RNAi^ (mCD8GFP, green). Follicle cells expressing Par-1^RNAi^ exhibits increased TR-avidin signal compared to nearby wild type cells. Scale bar denotes 50 μm. **(D-G)** Par-1 negatively regulates Fas2 display on the lateral side. **(D-E)** Superficial view of Stage 5/6 egg chamber expressing Par-1^RNAi^, marked by mCD8GFP expression (green), shows premature Fas2 (magenta) removal. **(F-G)** A Cross-sectional image of anterior end of Stage 6/7 egg chamber expressing Par-1^RNAi^ (green) shows premature Fas2 removal from the from the lateral side of anterior follicle cells. **(H)** Par-1 physically interacts with Fas2. Par-1:GFP lysates immunoprecipitated with GFP antibodies (DSHB:12A6) show the presence of Fas2 (DSHB:1D4) in the indicated lanes. Same blot as in (H) stripped and re-probed with GFP antibody (DSHB:N86/8) to show the presence of precipitated Par-1:GFP in the indicated lanes. **(I)** Genetic reduction of Par-1 rescues the morphogenesis defects observed upon TOR depletion. Compare cell area of indicated genotypes. Error bars indicate SEM. **** denotes p-value < 0.0001 **(J-L’’’)** Premature Fas2 (green or black) removal phenotype observed upon Par-1 Depletion (Stinger.nls, magenta) can be rescued by genetic reduction of Hsc70-4, the clathrin uncoating ATPase. **(J-J’’’)** Fas2 removal phenotype in *hsc70-4^L3929^*/+. No instances of premature Fas2 removal were observed in this background. **(K-K’’’)** Fas2 removal phenotype in Par-1^RNAi^ expressing follicle cells. **(L-L’’’)** Fas2 removal phenotype in Par-1^RNAi^ + *hsc70-4^L3929^/+* background.

### Par-1 physically interacts with Fas2

Given that our previous observations suggest that Par-1 negatively regulates endocytosis and also promotes membrane retention of Fas2 in cuboidal cells, we next asked how Par-1 might be regulating Fas2 removal in the AFCs. Immunostaining of Egg chambers with Par-1 (Par-1:GFP) and Fas2 exhibit some overlap in the expression pattern on the lateral side of developing follicle cells (Fig S7B-B’’). This motivated us to test whether Par-1 physically interacts with Fas2. To test this, we utilized a Par-1:GFP protein trap line to immunoprecipitate Par-1 from ovarian lysates using anti-GFP mAb, and immunoblotted for Fas2 (1D4 mAb) ^51^. Immunoprecipitation of Par-1:GFP lysates with a GFP specific IgG but not control IgG, revealed the presence of Fas2 in the same immunocomplex containing Par-1. Consistent with the immunolocalization results, we found evidence that Par-1 can physically interact with Fas2 in the fly ovaries (Fig 8H, Full blot shown in Fig S7F).

As Par-1 and Fas2 can physically interact, we specifically asked if Par-1 was modulating Fas2 levels in the AFCs through endocytosis. To test the possibility, we conducted a genetic interaction between Par-1 and Hsc70-4, the clathrin uncoating ATPase. We reasoned that if Par-1 depletion promotes premature Fas2 removal, then perturbing endocytosis in this background should dampen the premature Fas2 removal phenotype. To do this, we generated Par-1^RNAi^ expressing clones in follicle cells and quantified the instances of premature Fas2 removal (between stages 5-8). We similarly assessed the phenotype in Par-1 knockdown in the background of the Hsc70-4 null allele, *hsc70-4^L3929,^* and the heterozygous *hsc70-4^L3929^/+* alone ^43, 52^. We found, roughly 92% of egg chambers in Par-1^RNAi^ (n=83) displayed premature loss of Fas2, compared to 72% of egg chambers in Par-1^RNAi^ with a *hsc70-4^L3929^* allele (n=72), while none was observed in *hsc70-4^L3929/+^* alone (n=76). Hence, this genetic interaction between Par-1 and Hsc70-4 suggests that the premature Fas2 removal observed in the Par-1^RNAi^ background is dependent on endocytosis (Fig 8J-L’’’).

Altogether our results above suggest that Par-1 physically interacts with Fas2 and functions through endocytosis to modulate Fas2 removal in the shape transitioning AFCs.

## Discussion

Cell shape transformation during epithelial morphogenesis forms an integral component of metazoan development and survival. Here we have evaluated the role of TOR signalling in mediating epithelial cell shape change specifically focusing on how a cuboidal cell transforms into a squamous shape. By employing the model of AFCs in the vitellogenic *Drosophila* egg chambers, we demonstrate that growth promoting TOR signalling mediates the shape transformation of cuboidal AFCs to squamous fate. Our results suggest that TORC1 negatively regulates the serine-threonine polarity kinase Par-1 to facilitate the removal of Fas2 protein from the lateral membrane of the AFCs and assist in the shape transformation of cuboidal follicle cells to squamous fate.

### Role of TOR in epithelial morphogenesis

The role of TOR signalling is well-defined in mediating cell growth and proliferation, in several eukaryotic organisms. In addition, it has been shown that TOR signalling regulates morphogenesis in several model systems like *C. elegans, Drosophila,* Zebrafish, and mice potentiating early embryogenesis, aiding in transitioning through intermediate developmental stages, and facilitating differentiation of organs like skin, intestine, etc ^53^. Incidentally, Makky et al reported in 2007 that Rapamycin treatment inhibited differentiation of intestinal cell progenitor and stalled them in the cuboidal shape configuration itself ^54^. Though there are indications in the literature that TOR may affect cell shape transition in developing organs, there is hardly any study that investigates this aspect of TOR signalling in metazoan development.

From our work, several main conclusions can be drawn. First, we provide evidence that TOR signaling regulates cell shape change during epithelial morphogenesis. Second, unlike the known effectors of TOR, we demonstrate that Par-1 plays a significant role downstream of TORC1 to modulate the Fas2 removal from the lateral membrane. Though the cuboidal to squamous transition is associated with expansive growth of the follicle cells, intriguingly our results suggest that it is independent of the growth-promoting (S6 kinase) arm of TOR signaling as downregulation of S6 kinase didn’t affect AFCs. As over-expression of REPTOR^CA^ doesn’t impede morphogenesis associated with AFC, we believe that the transcriptional component of TORC1 that functions through modulation of REPTOR activity doesn’t play any significant role in mediating this shape change. Third, we believe that the epithelial morphogenesis associated with AFCs is not linked to the cytoskeleton as we didn’t observe any conspicuous difference in the levels of tubulin between the control and TOR-depleted anterior follicle cells. Fourth, since TOR depletion didn’t perturb the levels of Par-1, it suggests that TORC1 modulates Par-1 post-translationally to regulate Fas2 endocytosis.

Now the question arises as to how TOR signaling is mediating Fas2 removal. Our results suggest that TORC1 negatively regulates Par-1 to modulate Fas2 removal in the transitioning AFCs. We demonstrate that Par-1 negatively regulates endocytosis in AFCs thus impeding Fas2 removal in shape-transitioning AFCs. Though our data suggest that Par-1 and Fas2 physically interact, we don’t think that Par-1’s presence on the lateral membrane physically impedes the Fas2 removal. The reason for this assumption is that under WT conditions, we didn’t observe any dip in Par-1 levels before Fas2 removal from the lateral membrane of the AFCs. However, we can’t rule out the possibility of local spatial changes or alteration in the activity of Par-1 that may favor the Fas2 removal in response to TOR signaling.

In the context of other players mediating this shape change, we know that serine-threonine kinase Tao also mediates Fas2 removal from the lateral membrane of AFCs. It will be worth investigating if TOR and Tao function together to synchronize the shape change of AFCs. On another note, unlike the report of Gomez et al, we did observe that serine-threonine Par-1 kinase impedes Fas2 removal from AFCs. We believe that this difference in observation may be attributed to the use of different Par-1^RNAi^ constructs in the earlier study which was targeting only a subset of Par-1 isoforms^27^. Nevertheless, it would be worth testing if TOR functions through Tao to downregulate Par-1 to mediate Fas2 endocytosis in AFCs.

Though our understanding of TOR signaling has come a long way since the discovery of Rapamycin around three decades ago, of late we are slowly beginning to understand how TOR signaling regulates metazoan morphogenesis. Given that cuboidal to squamous cell shape transition is associated with the organogenesis of skin, lungs, and kidneys, it would be worth examining if the epithelial morphogenesis events observed in the above organs follow a similar molecular mechanism as reported in *Drosophila* AFCs.

## Materials and Methods

All stocks and crosses were maintained at 25°C unless otherwise stated. Experiments involving the expression of RNAi and overexpression constructs of transgenes were fattened at 29 °C for 20-24 hours. For live imaging and in the interaction between *TOR^RNAi^* and *hsc70-4^L3929^*, the fattening was done at 25 °C instead of 29 °C. For temperature-sensitive Shi transgene, the flies were fattened at 31 °C. For GR1-Gal4 experiments, the eclosed flies were shifted to 16 °C for 2-4 days followed by fattening at 29 °C for dissection. Induction of transgenes using Flipout/AyGal4 was induced by heat shocking at 37 °C 3 times for a day followed by dissection after 4 days. Clone generation for endocytosis screening and *TOR^ΔP^* mutant clones was done according to Sharma et al., 2017, followed by fattening at 29 °C and 25 °C, respectively ^55^.

### Quantification of Cell Area

Area information was acquired by outlining cell perimeters in Zen Blue software. The cell perimeter was judged using the GFP marker expression in the clones and/or with armadillo or tubulin co-staining. For cuboidal cells, stage 7 was considered and for squamous cells stage 10 was considered.

### Quantification of Cell Volume

ZenBlue software was used for extracting cell features – cell area and cell height. Cell area was obtained similarly as described previously, using the Armadillo channel. For cell height, the extent of the Armadillo signal across the number of confocal sections was obtained by multiplying z-steps by the z-step size (0.44 μm). Finally, the area was multiplied by the calculated height to obtain the volume for each specific cell.

### Immunostaining

Ovaries were dissected in Schneider’s *Drosophila* medium supplemented with 10 % fetal bovine serum. Fixation was done for 20 minutes at RT, followed by a wash using PBS + 0.3% Triton X-100 (PT_0.3_). Samples were blocked using 0.5% BSA in PT_0.3_ for 1 hour followed by primary antibody incubation O/N. After primary antibody incubation, samples were washed four times, followed by appropriate secondary antibody (Life Technologies, 1:500 dilution), and incubated for 1.5 hours. This was followed by four washes, including nuclear labeling with DAPI, and mounted in Vectashield H-1000. For Fas2 staining, PT_0.3_ was substituted with PT_0.1_ and washed with PBS. For pS6 staining, ovaries were directly pulled into 4% PFA solution, fixed for 15 minutes and staining continued as indicated above, using PT_0.1_.

### Coverslip and Confocal Dish preparation for Live Imaging

Coverslips were cleaned with 70% EtOH and affixed onto the bottom surface of the tissue culture dish, using vacuum silicone grease. Then, spotted with Poly-D Lysine (PDL, Cat# P7280 Sigma) solution (100µg/ml) in water, at the center of the coverslip, followed by microwaving for 15 seconds, removal of excess fluid and drying O/N in 37°C incubators. Coated coverslips were UV treated for 15 minutes before use. Excess/unbound PDL was washed off from the coverslip with a brief incubation and washed with the live imaging media, before applying the egg chambers,

### Live imaging of Cell Shape Transition

Live imaging was performed essentially as described in Prasad et al., 2007 with a few modifications ^56^. Live imaging media was supplemented with 20-hydroxyEcdysone (10mM 2000X stock in DMSO). A tissue culture dish instead of lummox apparatus, prepared as described above, was used. Excess/unbound PDL was washed off from the coverslip with a brief incubation and washed with live imaging media, before applying the egg chambers. Tissue paper soaked in water was kept inside the dish and covered with a lid, before mounting it on the microscope.

### Live imaging setup, processing and analysis

Live imaging was performed on a Leica SP8 laser scanning confocal in Lightning mode. Samples were imaged with a 40X objective, 5 minutes apart, in bidirectional scan mode, keeping resolution to 89 nm, with a z-step size of 1 μm, scan speed of 400-600, pinhole of 1.2 Airy units using a HyD detector. Images were taken using both single and multi-position acquisition. The stack was refocused and revised from time to time.

Post-imaging, Maximum Z projection of each stack over time was exported using Leica LasX software and assembled in Fiji to generate movies or montages.

In Fiji, after scale calibration, sub-anterior most cells, roughly 5-10 in number, were outlined and tracked overtime. The mean area of these outlined cells was obtained for analysis of cell area over time. We limited our analysis to time frames after rotation of egg chambers had stopped. For calculating % increase in over 1^st^ frame, the % increase in cell area over first frame was calculated for each time frame, averaged across samples and plotted. Calculation of the mean rate of expansion was done as follows - the fractional increase in cell area of each time point over first frame was obtained and divided by the time elapsed in minutes.

### Texas Red conjugated Avidin Internalisation Assay and Quantification

Briefly, 2-3 ovaries of experimental genotype were dissected in room temperature Live imaging media with Hoechst 33342. Individual egg chambers were teased out from the ovarian sheath using fine forceps. Live imaging media was gently replaced with TR-avidin dye (200µg/ml) and incubated at 29°C for 30minutes, with intermittent mixing. Post incubation, egg chambers were rinsed with pre-chilled Live imaging media three times for 2-3 minutes, while on ice. Post-washing samples were mounted and imaged within 30 minutes. Egg chambers with very high Hoechst staining were not considered, as it indicated damaged cells with excess TR-Avidin signal. Images were acquired using Zeiss Apotome 2.0 with binning 2 to lower exposure time and facilitate faster imaging.

For quantification of internalised TR-avidin dye, intensity values of TR-avidin were generated using ZenBlue, by drawing an ROI or Region of Interest on the clonal cells, followed by copy pasting of same ROI into a region of non-clonal population, to keep same area of ROI. Then, TR-avidin intensity in clone was normalised to intensity of nearby control cells to obtain data points.

### Line profiling analysis of Par-1 GFP across Basal to Apical Membrane

Analysis was done on maximum intensity projections of cells beyond 4 cells from the polar cells, using Zen Zeiss Blue. Line profile tool was drawn starting from baso-lateral region to apico-lateral region, spanning roughly 3.5-4 μm, to obtain intensity profiles and plotted.

### Immunoprecipitation and Western Blotting

Around 30 flies were dissected in cold 0.2% CHAPS based lysis buffer LB (10mM Tris-Cl pH 7.4, 150mM NaCl, 0.2% CHAPS, 1X Protease Inhibitor Cocktail, 1X PhosSTOP phosphatase inhibitor cocktail, 1mM EDTA) and homogenized using a micropestle. Post homogenization, pipetting and vortexing was done before clarified at 16000g/4C for 15 minutes. Meanwhile, Protein G PLUS Agarose beads (# BB-PG001PB) beads were equilibrated using LB. The clarified lysate was transferred to a fresh tube and quantified using Bradford assay. The lysate was pre-cleared with 10 μl of equilibrated beads. A small aliquot was separated and stored on ice as Input. Around 200μg pre-cleared lysate was incubated with 1 ug Control IgG (Cat #sc-2002) and 1 ug anti-GFP (DSHB 12A6, BioReactor Supernatant), for 2-4 hours in a rotatory shaker at 4°C, followed by addition of 20ul of Beads and O/N incubation. Next day, the beads were washed with LB, boiled in 2x Lammeli buffer and loaded onto 6% SDS-PAGE gel and transferred to PVDF membrane for western blotting.

The blots were probed with 1D4 anti-Fas2 (1:250, DSHB), followed by stripping and re-probing with N86/8 anti-GFP (1:300, DSHB). Blocking and antibody incubations were done in presence of 5% Skim Milk in washing solution composed of TBS-Tween 20 (0.05%).

### RNA isolation, cDNA Synthesis and semi-qRT-PCR

RNA was isolated from 4-5 pairs of ovaries with TRIzol reagent (Invitrogen Catalogue No. 15596026) and cDNA was synthesized from 1µg of RNA, using RevertAid Reverse Transcriptase (Catalogue No.-Thermo Scientific EP0441) following manufacturer instructions. For *rictor* and *rp49* semi qPCR, 1.0 μl and 0.25 μl of undiluted cDNA was used as a template for a 25 cycle and 20 cycle reactions, respectively. Full reaction was loaded for analysis.

Primers Sequences:

rictor forward – GCGTCACCTCCATAACCCG and

rictor reverse – ACCGCAGATGTTCCTCGTTTG (primer pair PP35665, FlyPrimer Bank,^57^) and

rp49 forward: CTAAGCTGTCGCACAAATGGC, rp49 reverse: AACTTCTTGAATCCGGTGGGC.

### Statistical tests

Difference of means determined by Students’ t-test with Welch’s Correction in GraphPad Prism 6.0. All error bars indicate Standard Deviation of Mean. All figures utilize, the following range for assigning significance of means: p-value <0.0001 is designated as ****, p-value <0.001 is designated as ***, p-value <0.01 is designated as **, 0.05> p >0.01 is designated as * and >0.05 as ns or not significant.

## Supporting information

Movie 1 Control

Movie 2 TORRNAi

## Supplementary Figures

**Supplementary Figure 1:**
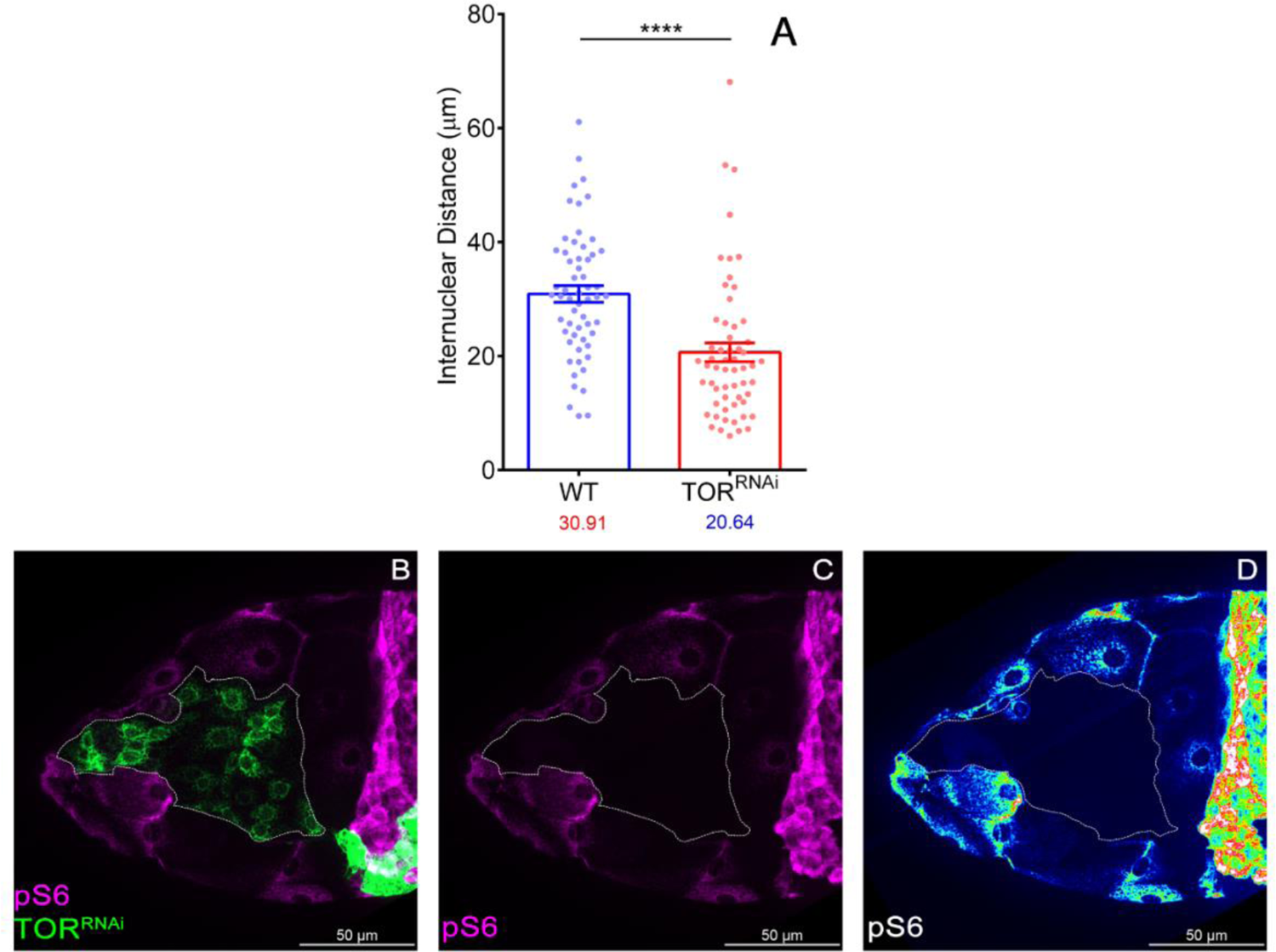
Validation of TOR^RNAi^ using TORC1 reporter, pS6 and Nuclei Clustering in TOR^RNAi^. **(A)** Comparison of internuclear distance in nearby control cells and TOR^RNAi^ expressing anterior follicle cells clones. **(B-D)** Validation of TOR^RNAi^ using pS6 antibody. Anterior cells expressing TOR^RNAi^ are marked by mCD8GFP expression and devoid of pS6 staining. Error bars indicate SEM. **** indicates p-value < 0.0001 in Students’ t-test with Welch’s Correction

**Supplementary Figure 2:**
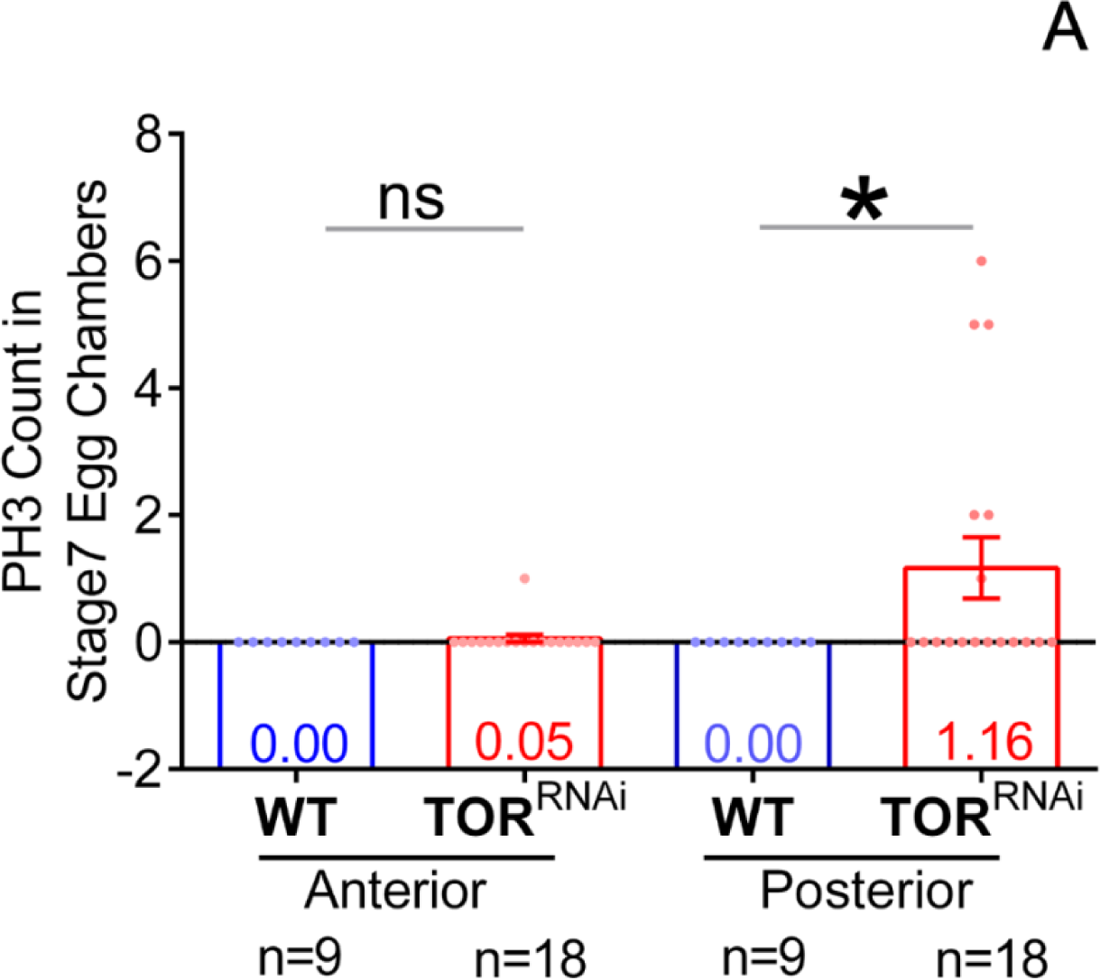
Status of Mitosis in TOR Depleted follicle cells. **(A)** TOR depletion leads to sustained PH3 staining in the posterior follicle cells till Stage 7. PH3 count is significantly different in the posterior side. Error bars indicate SEM.* indicates p-value < 0.05 in Students’ t-test with Welch’s correction.

**Supplementary Figure 3:**
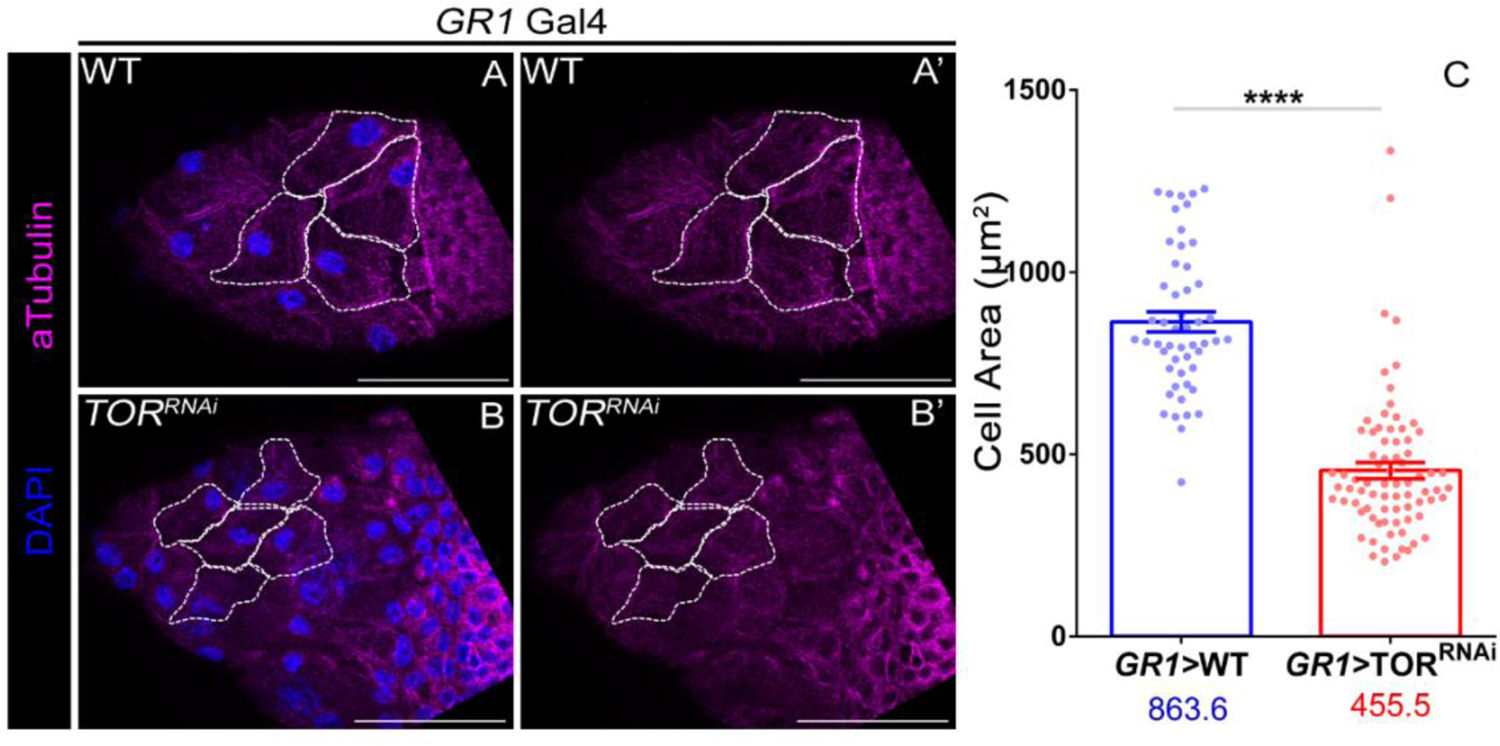
*GR1-Gal4* specific knockdown of TOR in squamous cell morphogenesis. **(A-B)** Comparison of Cell Area in Control **(A-A’)** and TOR^RNAi^ **(B-B’)** in background of follicle specific *GR1*-Gal4 driver. **(C)** Quantification of cell area in the indicated backgrounds. Error bars indicate SEM. **** indicate p-value < 0.0001, in Students’ t-test with Welch’s correction.

**Supplementary Figure 4:**
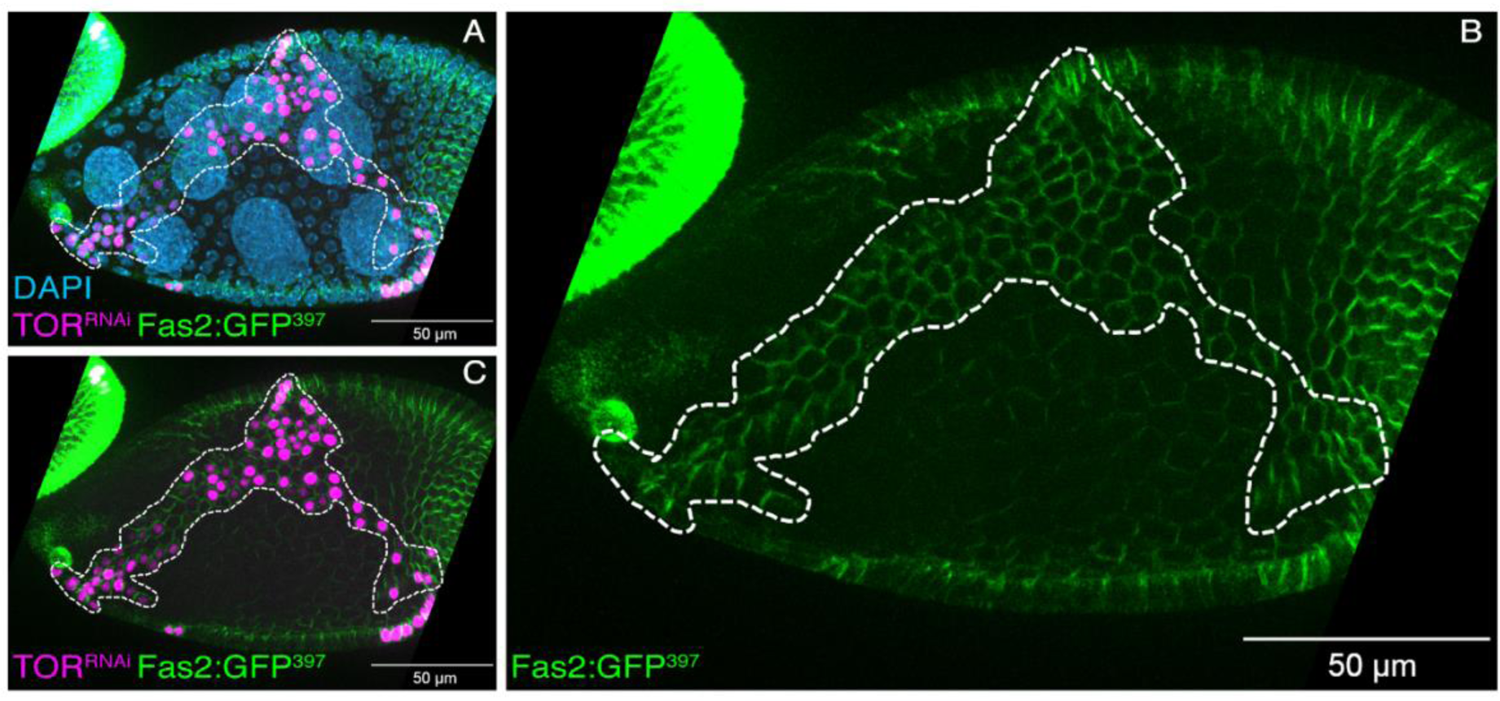
Delayed removal of endogenously tagged Fas2 in TOR depleted follicle cells. **(A-C)** Delayed Fas2:GFP^397^(green) removal in the TOR depleted follicle cells indicated by expression of Stinger.nls (magenta)

**Supplementary Figure 5:**
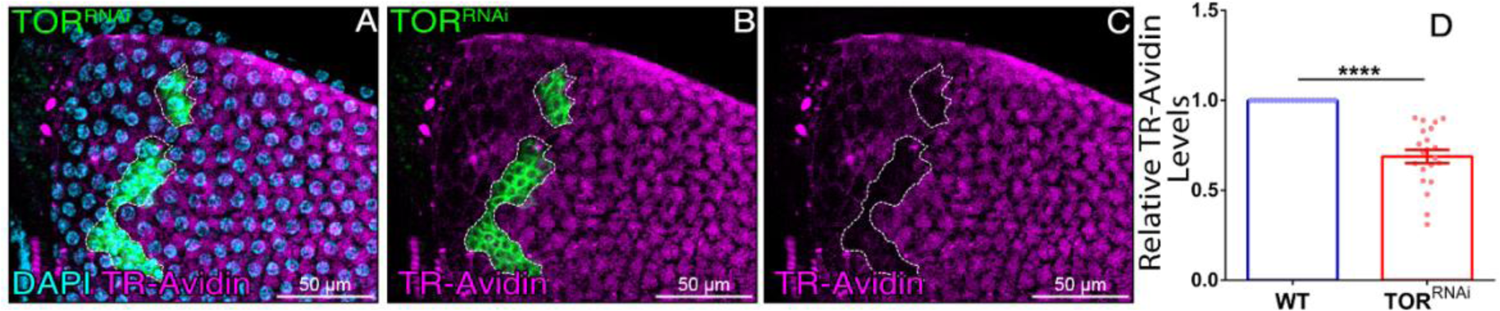
TOR depletion results in reduced endocytosis. **(A-C)** TOR depleted follicle cells (green), exhibit reduced internalization of TexasRed conjugated to Avidin or TR-Avidin (magenta) and Hoechst labelling nuclei in live cells (cyan). **(D)** Plot showing quantification of TR-avidin uptake in TOR^RNAi^ clones and nearby wild type cells. Error bars indicate SEM. **** indicate p-value < 0.0001, in Students’ t-test with Welch’s correction.

**Supplementary Figure 6:**
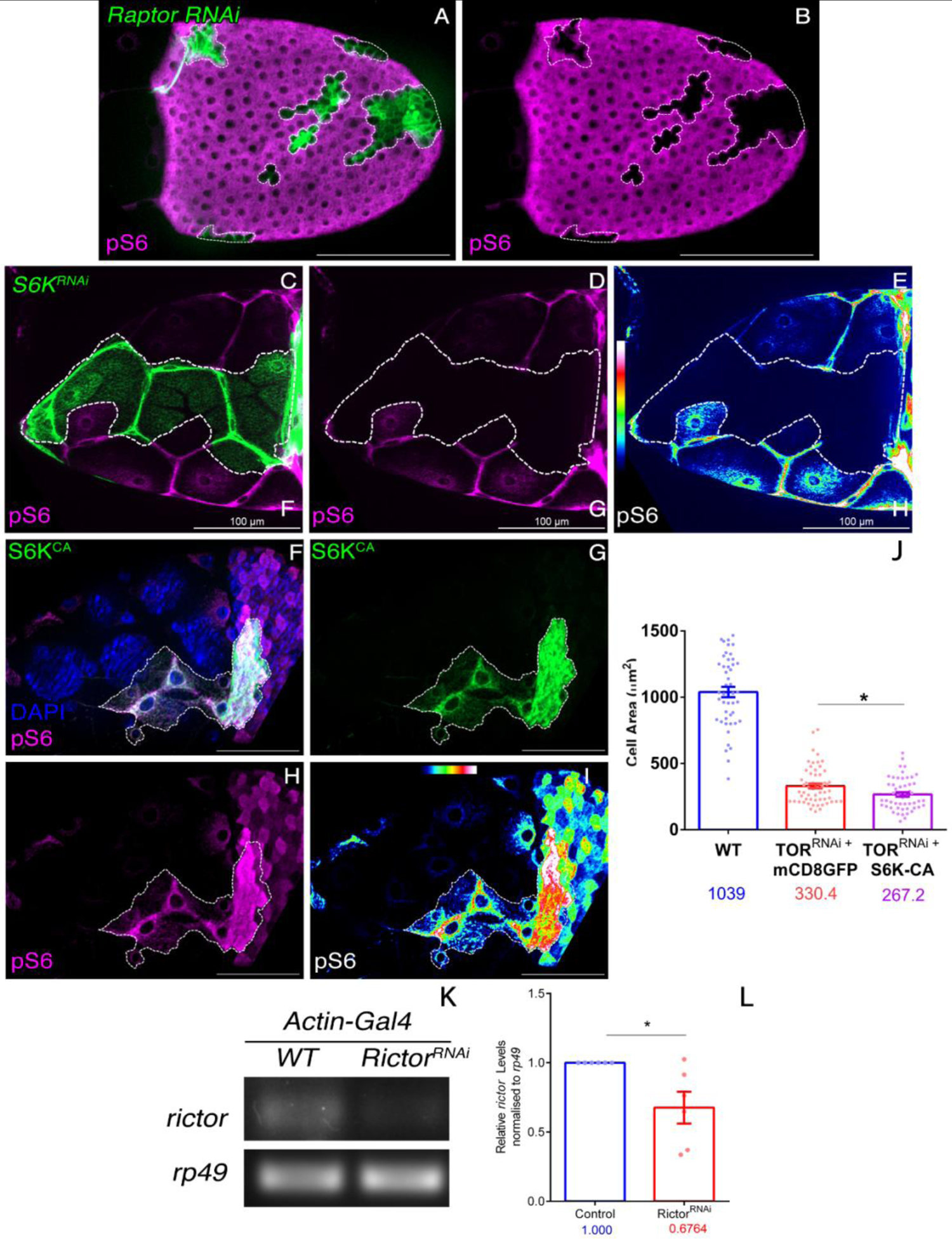
Validation of constructs and lack of rescue of TOR^RNAi^ phenotypes by S6K-CA overexpression. **(A-B)** Validation of Raptor^RNAi^ (mCD8GFP, green) by pS6 antibody (magenta). **(C-E)** Validation of S6K^RNAi^ (mCD8GFP, green) by pS6 antibody (magenta and heatmap) **(F-I)** Validation of S6K^CA^ (mCD8GFP, green) by pS6 antibody (magenta and heatmap) **(J)** Cell size defects of TOR^RNAi^ are not rescued by S6K-CA overexpression. **(K-L)** Validation of Rictor^RNAi^ construct using semi-quantitative RT-PCR, target gene: *rictor* normalised against *rp49* Error bars denote SEM. * Represent p-value <0.05 in Students’ t-test with Welch’s correction.

**Supplementary Figure 7:**
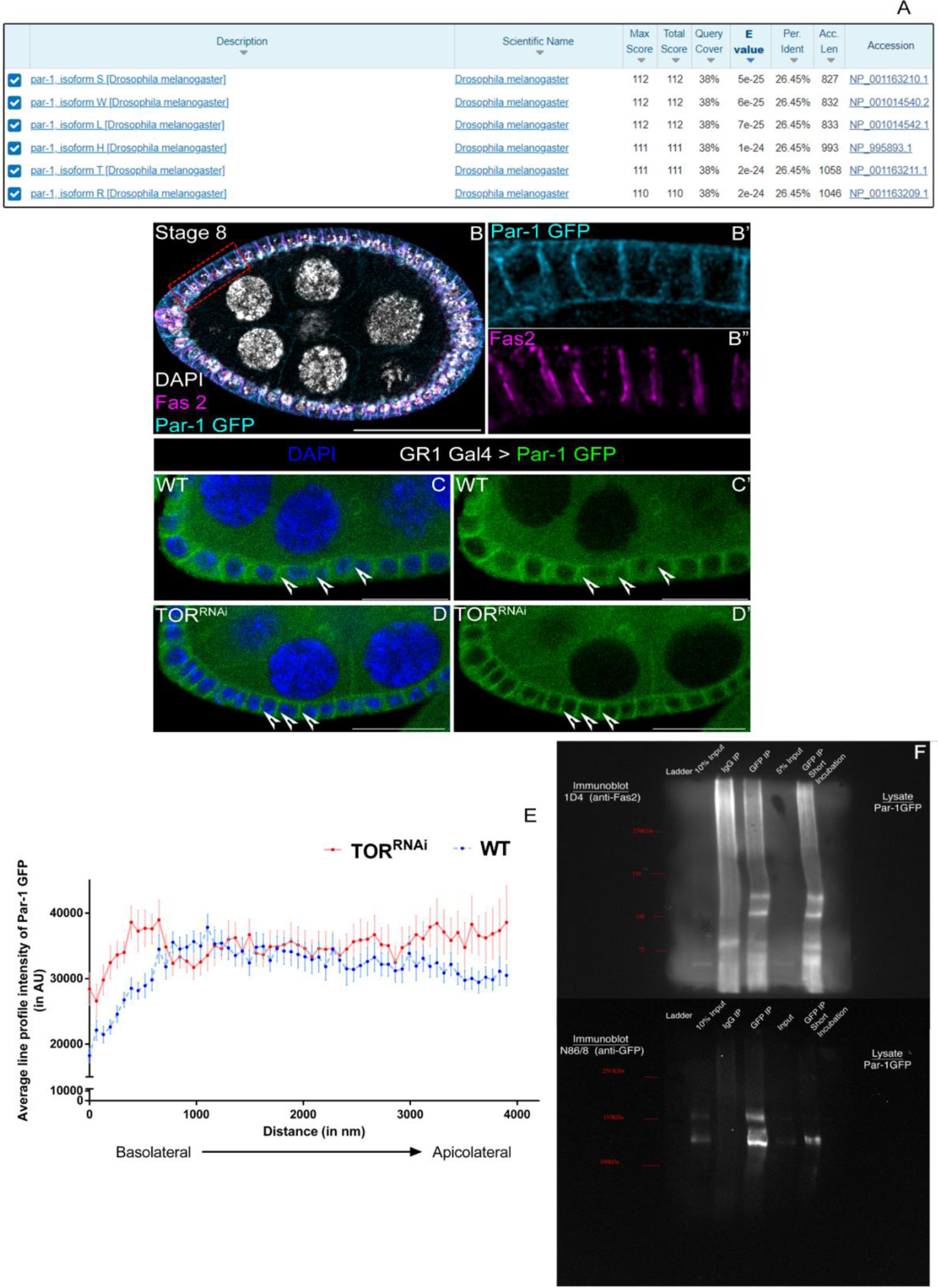
Similarity of *Dmel* Par-1 to *Scer* Npr1, colocalization of Par-1 and Fas2 in egg chambers, status of Par-1 in TOR knockdown conditions and physical interaction of Par-1 with Fas2. **(A)** Protein BLAST analysis of yeast Npr1 protein sequence (Uniprot ID P22211) against *Drosophila melanogaster* sequences, reveal Par-1 as the closest homolog. **(B-B’’)** Comparison of Fas2 (magenta) and endogenous Par-1:GFP (αGFP, cyan) localisation in egg chambers from homozygous Par-1:GFP flies (Carnegie Protein Trap CC01981). **(C-D)** Comparison of Par-1:GFP (endogenous signal, green) levels in control (C-C’) and TOR^RNAi^ (D-D’), expressing egg chambers. **(E)** Plot showing the comparison of line profiles (in AU) of Par-1:GFP in Control and TOR^RNAi^ egg chambers, drawn across basal to apical side (in nm). Note, the altered distribution of Par-1:GFP signal across the basolateral to apicolateral domain under TOR knockdown compared to the control. Error bars indicate SEM. **(F)** Full Blots showing the physical interaction between Par-1 and Fas2. Ovaries from homozygous Par-1:GFP flies were pulled down using 12A6 αGFP antibody (bioreactor supernatant), and probed with aFas2 (1D4) antibody, stripped and reprobed with N86/8 αGFP antibody.

---

**Movie 1: Visualising squamous cell morphogenesis in control egg chambers labelled with DE-cadherin:GFP** Time lapse images of control egg chambers obtained after maximum intensity projection of each time point. Genotype: E-Cadherin:GFP; *GR1*-Gal4*/+*.

**Movie 2: Visualising squamous cell morphogenesis in TOR depleted egg chambers visualised with DE-cadherin:GFP** Time lapse images of TOR depleted egg chambers obtained after maximum intensity projection of each time point. Genotype: E-cadherin:GFP; *GR1*-Gal4/TOR^RNAi^.

## Notes

### Competing Interest Statement

The authors have declared no competing interest.

## References

1. Gary C. Schoenwolf & Steven B. Bleyl & Philip R. Brauer & Philippa H. Francis-West (2021). Larsen’s Human Embryology, ISBN Number-9780323696043 6th Edition.

2. Hu, M.S., Borrelli, M.R., Hong, W.X., Malhotra, S., Cheung, A.T.M., Ransom, R.C., Rennert, R.C., Morrison, S.D., Lorenz, H.P., and Longaker, M.T. (2018). Embryonic skin development and repair. https://doi.org/10.1080/15476278.2017.1421882 14, 46–63. 10.1080/15476278.2017.1421882.

3. Schell, C., Wanner, N., and Huber, T.B. (2014). Glomerular development - Shaping the multi-cellular filtration unit. Semin Cell Dev Biol 36, 39–49. 10.1016/J.SEMCDB.2014.07.016.

4. Bertet, C., Sulak, L., and Lecuit, T. (2004). Myosin-dependent junction remodelling controls planar cell intercalation and axis elongation. Nature 429, 667–671. 10.1038/nature02590.

5. Blankenship, J.T., Backovic, S.T., Sanny, J.S.S.P., Weitz, O., and Zallen, J.A. (2006). Multicellular Rosette Formation Links Planar Cell Polarity to Tissue Morphogenesis. Dev Cell 11, 459–470. 10.1016/j.devcel.2006.09.007.

6. Harris, T.J.C., and Peifer, M. (2007). aPKC Controls Microtubule Organization to Balance Adherens Junction Symmetry and Planar Polarity during Development. Dev Cell 12, 727–738. 10.1016/j.devcel.2007.02.011.

7. Brigaud, I., Duteyrat, J.L., Chlasta, J., le Bail, S., Couderc, J.L., and Grammont, M. (2015). Transforming growth factor b/activin signalling induces epithelial cell flattening during drosophila oogenesis. Biol Open 4, 345–354. 10.1242/bio.201410785.

8. Kuo, Y., Huang, H., Cai, T., and Wang, T. (2015). Target of rapamycin complex 2 regulates cell growth via Myc in drosophila. Sci Rep 5, 1–10. 10.1038/srep10339.

9. Laplante, M., and Sabatini, D.M. (2012). MTOR signaling in growth control and disease. Cell 149, 274–293. 10.1016/j.cell.2012.03.017.

10. Wullschleger, S., Loewith, R., and Hall, M.N. (2006). TOR signaling in growth and metabolism. Cell 124, 471–484. 10.1016/j.cell.2006.01.016.

11. Loewith, R., Jacinto, E., Wullschleger, S., Lorberg, A., Crespo, J.L., Bonenfant, D., Oppliger, W., Jenoe, P., and Hall, M.N. (2002). Two TOR complexes, only one of which is rapamycin sensitive, have distinct roles in cell growth control. Mol Cell 10, 457–468. 10.1016/S1097-2765(02)00636-6.

12. Kunz, J., Henriquez, R., Schneider, U., Deuter-Reinhard, M., Movva, N.R., and Hall, M.N. (1993). Target of rapamycin in yeast, TOR2, is an essential phosphatidylinositol kinase homolog required for G1 progression. Cell 73, 585– 596. 10.1016/0092-8674(93)90144-F.

13. Burnett, P.E., Barrow, R.K., Cohen, N.A., Snyder, S.H., and Sabatini, D.M. (1998). RAFT1 phosphorylation of the translational regulators p70 S6 kinase and 4E-BP1. Proc Natl Acad Sci U S A 95, 1432–1437. 10.1073/PNAS.95.4.1432/ASSET/BC13850A-4329-4DC5-80B3-166C31491D73/ASSETS/GRAPHIC/PQ0384115005.JPEG.

14. Isotani, S., Hara, K., Tokunaga, C., Inoue, H., Avruch, J., and Yonezawa, K. (1999). Immunopurified Mammalian Target of Rapamycin Phosphorylates and Activates p70 S6 Kinase α in Vitro. Journal of Biological Chemistry 274, 34493– 34498. 10.1074/JBC.274.48.34493.

15. Moreno-Torres, M., Jaquenoud, M., Péli-Gulli, M.P., Nicastro, R., and De Virgilio, C. (2017). TORC1 coordinates the conversion of Sic1 from a target to an inhibitor of cyclin-CDK-Cks1. Cell Discov 3, 1–12. 10.1038/celldisc.2017.12.

16. Balcazar, N., Sathyamurthy, A., Elghazi, L., Gould, A., Weiss, A., Shiojima, I., Walsh, K., and Bernal-Mizrachi, E. (2009). mTORC1 activation regulates β-cell mass and proliferation by modulation of cyclin D2 synthesis and stability. Journal of Biological Chemistry 284, 7832–7842. 10.1074/jbc.M807458200.

17. García-Martínez, J.M., and Alessi, D.R. (2008). mTOR complex 2 (mTORC2) controls hydrophobic motif phosphorylation and activation of serum- and glucocorticoid-induced protein kinase 1 (SGK1). Biochemical Journal 416, 375– 385. 10.1042/BJ20081668.

18. Yuan, M., Pino, E., Wu, L., Kacergis, M., and Soukas, A.A. (2012). Identification of Akt-independent regulation of hepatic lipogenesis by mammalian target of rapamycin (mTOR) complex 2. Journal of Biological Chemistry 287, 29579– 29588. 10.1074/jbc.M112.386854.

19. Oh, W.J., and Jacinto, E. (2011). mTOR complex 2 signaling and functions. Cell Cycle 10, 2305–2316. 10.4161/cc.10.14.16586.

20. Datta, S.R., Dudek, H., Xu, T., Masters, S., Haian, F., Gotoh, Y., and Greenberg, M.E. (1997). Akt phosphorylation of BAD couples survival signals to the cell-intrinsic death machinery. Cell 91, 231–241. 10.1016/S0092-8674(00)80405-5.

21. Horne-Badovinac, S., and Bilder, D. (2005). Mass transit: Epithelial morphogenesis in the Drosophila egg chamber. Developmental Dynamics 232, 559–574. 10.1002/dvdy.20286.

22. Kolahi, K.S., White, P.F., Shreter, D.M., Classen, A.K., Bilder, D., and Mofrad, M.R.K. (2009). Quantitative analysis of epithelial morphogenesis in Drosophila oogenesis: New insights based on morphometric analysis and mechanical modeling. Dev Biol 331, 129–139. 10.1016/j.ydbio.2009.04.028.

23. Montell, D.J. (2003). Border-cell migration: The race is on. Nat Rev Mol Cell Biol 4, 13–24. 10.1038/nrm1006.

24. Wu, X., Tanwar, P.S., and Raftery, L.A. (2008). Drosophila follicle cells: Morphogenesis in an eggshell. Semin Cell Dev Biol 19, 271–282. 10.1016/j.semcdb.2008.01.004.

25. Grammont, M. (2007). Adherens junction remodeling by the Notch pathway in Drosophila melanogaster oogenesis. Journal of Cell Biology 177, 139–150. 10.1083/jcb.200609079.

26. Sahu, A., Karmakar, S., Halder, S., Ghosh, G., Acharjee, S., Dasgupta, P., Ghosh, R., Deshpande, G., and Prasad, M. (2021). Germline soma communication mediated by gap junction proteins regulates epithelial morphogenesis. PLoS Genet 17, e1009685. 10.1371/JOURNAL.PGEN.1009685.

27. Gomez, J.M., Wang, Y., and Riechmann, V. (2012). Tao controls epithelial morphogenesis by promoting fasciclin 2 endocytosis. Journal of Cell Biology 199, 1131–1143. 10.1083/jcb.201207150.

28. LaFever, L., Feoktistov, A., Hsu, H.J., and Drummond-Barbosa, D. (2010). Specific roles of Target of rapamycin in the control of stem cells and their progeny in the Drosophila ovary. Development 137, 2117–2126. 10.1242/dev.050351.

29. Kim, W., Jang, Y.G., Yang, J., and Chung, J. (2017). Spatial Activation of TORC1 Is Regulated by Hedgehog and E2F1 Signaling in the Drosophila Eye. Dev Cell 42, 363–375.e4. 10.1016/j.devcel.2017.07.020.

30. Romero-Pozuelo, J., Demetriades, C., Schroeder, P., and Teleman, A.A. (2017). CycD/Cdk4 and Discontinuities in Dpp Signaling Activate TORC1 in the Drosophila Wing Disc. Dev Cell 42, 376–387.e5. 10.1016/j.devcel.2017.07.019.

31. Zhang, H., Stallock, J.P., Ng, J.C., Reinhard, C., and Neufeld, T.P. (2000). Regulation of cellular growth by the Drosophila target of rapamycin dTOR. Genes Dev 14, 2712–2724. 10.1101/gad.835000.

32. Lee, T., and Luo, L. (2001). Mosaic analysis with a repressible cell marker (MARCM) for Drosophila neural development. Trends Neurosci 24, 251–254. 10.1016/S0166-2236(00)01791-4.

33. Martín-Castellanos, C., and Edgar, B.A. (2002). A characterization of the effects of Dpp signaling on cell growth and proliferation in the Drosophila wing. Development 129, 1003–1013. 10.1242/DEV.129.4.1003.

34. Bai, J., and Montell, D. (2002). Eyes Absent, a key repressor of polar cell fate duringDrosophila oogenesis. Development 129, 5377–5388. 10.1242/DEV.00115.

35. Roth, S., Shira Neuman-Silberberg, F., Barcelo, G., and Schüpbach, T. (1995). cornichon and the EGF receptor signaling process are necessary for both anterior-posterior and dorsal-ventral pattern formation in Drosophila. Cell 81, 967–978. 10.1016/0092-8674(95)90016-0.

36. Huang, J., Zhou, W., Dong, W., Watson, A.M., and Hong, Y. (2009). Directed, efficient, and versatile modifications of the Drosophila genome by genomic engineering. Proc Natl Acad Sci U S A 106, 8284–8289. 10.1073/pnas.0900641106.

37. Haigo, S.L., and Bilder, D. (2011). Global tissue revolutions in a morphogenetic movement controlling elongation. Science (1979) 331, 1071–1074. 10.1126/science.1199424.

38. Laiouar, S., Berns, N., Brech, A., and Riechmann, V. (2020). RabX1 Organizes a Late Endosomal Compartment that Forms Tubular Connections to Lysosomes Consistent with a “Kiss and Run” Mechanism. Current Biology 30, 1177–1188.e5. 10.1016/j.cub.2020.01.048.

39. Silies, M., and Klämbt, C. (2010). APC/CFzr/Cdh1-dependent regulation of cell adhesion controls glial migration in the Drosophila PNS. Nat Neurosci 13, 1357– 1364. 10.1038/nn.2656.

40. Grenningloh, G., Jay Rehm, E., and Goodman, C.S. (1991). Genetic analysis of growth cone guidance in drosophila: Fasciclin II functions as a neuronal recognition molecule. Cell 67, 45–57. 10.1016/0092-8674(91)90571-F.

41. Ibar, C., Cataldo, V.F., Vásquez-Doorman, C., Olguín, P., and Glavic, Á. (2013). Drosophila p53-related protein kinase is required for PI3K/TOR pathway-dependent growth. Development (Cambridge) 140, 1282–1291. 10.1242/dev.086918.

42. Hennig, K.M., Colombani, J., and Neufeld, T.P. (2006). TOR coordinates bulk and targeted endocytosis in the Drosophila melanogaster fat body to regulate cell growth. Journal of Cell Biology 173, 963–974. 10.1083/jcb.200511140.

43. Chang, H.C., Newmyer, S.L., Hull, M.J., Ebersold, M., Schmid, S.L., and Mellman, I. (2002). Hsc70 is required for endocytosis and clathrin function in Drosophila. Journal of Cell Biology 159, 477–487. 10.1083/jcb.200205086.

44. Loewith, R., Jacinto, E., Wullschleger, S., Lorberg, A., Crespo, J.L., Bonenfant, D., Oppliger, W., Jenoe, P., and Hall, M.N. (2002). Two TOR complexes, only one of which is rapamycin sensitive, have distinct roles in cell growth control. Mol Cell 10, 457–468. 10.1016/S1097-2765(02)00636-6.

45. Wang, L., Lawrence, J.C., Sturgill, T.W., and Harris, T.E. (2009). Mammalian target of rapamycin complex 1 (mTORC1) activity is associated with phosphorylation of raptorby mTOR. Journal of Biological Chemistry 284, 14693– 14697. 10.1074/jbc.C109.002907.

46. Dos, D.S., Ali, S.M., Kim, D.H., Guertin, D.A., Latek, R.R., Erdjument-Bromage, H., Tempst, P., and Sabatini, D.M. (2004). Rictor, a novel binding partner of mTOR, defines a rapamycin-insensitive and raptor-independent pathway that regulates the cytoskeleton. Current Biology 14, 1296–1302. 10.1016/J.CUB.2004.06.054.

47. Tiebe, M., Lutz, M., de La Garza, A., Buechling, T., Boutros, M., and Teleman, A.A. (2015). REPTOR and REPTOR-BP Regulate Organismal Metabolism and Transcription Downstream of TORC1. Dev Cell 33, 272–284. 10.1016/J.DEVCEL.2015.03.013.

48. MacGurn, J.A., Hsu, P.C., Smolka, M.B., and Emr, S.D. (2011). TORC1 regulates endocytosis via npr1-mediated phosphoinhibition of a ubiquitin ligase adaptor. Cell 147, 1104–1117. 10.1016/j.cell.2011.09.054.

49. Elbert, M., Rossi, G., and Brennwald, P. (2005). The yeast Par-1 homologs Kin1 and Kin2 show genetic and physical interactions with components of the exocytic machinery. Mol Biol Cell 16, 532–549. 10.1091/mbc.E04-07-0549.

50. Drewes, G., and Nurse, P. (2003). The protein kinase kin1, the fission yeast orthologue of mammalian MARK/PAR-1, localises to new cell ends after mitosis and is important for bipolar growth. FEBS Lett 554, 45–49. 10.1016/S0014-5793(03)01080-9.

51. Buszczak, M., Paterno, S., Lighthouse, D., Bachman, J., Planck, J., Owen, S., Skora, A.D., Nystul, T.G., Ohlstein, B., Allen, A., et al. (2007). The Carnegie Protein Trap Library: A Versatile Tool for Drosophila Developmental Studies. Genetics 175, 1505. 10.1534/GENETICS.106.065961.

52. Hing, H.K., Bangalore, L., Sun, X., and Artavanis-Tsakonas, S. (1999). Mutations in the heatshock cognate 70 protein (hsc4) modulate Notch signaling. Eur J Cell Biol 78, 690–697. 10.1016/S0171-9335(99)80037-5.

53. Ding, X., Bloch, W., Iden, S., Rüegg, M.A., Hall, M.N., Leptin, M., Partridge, L., and Eming, S.A. (2016). mTORC1 and mTORC2 regulate skin morphogenesis and epidermal barrier formation. Nature Communications 2016 7:17, 1–15. 10.1038/ncomms13226.

54. Makky, K., Tekiela, J., and Mayer, A.N. (2007). Target of rapamycin (TOR) signaling controls epithelial morphogenesis in the vertebrate intestine. Dev Biol 303, 501–513. 10.1016/J.YDBIO.2006.11.030.

55. Sharma, A., Halder, S., Felix, M., Nisaa, K., Deshpande, G., and Prasad, M. (2018). Insulin signaling modulates border cell movement in drosophila oogenesis. Development (Cambridge) 145. 10.1242/DEV.166165/VIDEO-3.

56. Prasad, M., Jang, A.C.C., Starz-Gaiano, M., Melani, M., and Montell, D.J. (2007). A protocol for culturing drosophila melanogaster stage 9 egg chambers for live imaging. Nat Protoc 2, 2467–2473. 10.1038/nprot.2007.363.

57. Hu, Y., Sopko, R., Foos, M., Kelley, C., Flockhart, I., Ammeux, N., Wang, X., Perkins, L., Perrimon, N., and Mohr, S.E. (2013). FlyPrimerBank: an online database for Drosophila melanogaster gene expression analysis and knockdown evaluation of RNAi reagents. G3 (Bethesda) 3, 1607–1616. 10.1534/g3.113.007021.

